# The sperm epigenome does not display recurrent epimutations in patients with severely impaired spermatogenesis

**DOI:** 10.1101/2020.02.10.941724

**Authors:** Elsa Leitão, Sara Di Persio, Sandra Laurentino, Marius Wöste, Martin Dugas, Sabine Kliesch, Nina Neuhaus, Bernhard Horsthemke

## Abstract

**Background:** In the past 15 years, numerous studies have described aberrant DNA methylation of imprinted genes (e.g. *MEST* and *H19*) in sperm of infertile patients, but the prevalence and genomic extent of abnormal methylation patterns have remained unknown.

**Results:** Using deep bisulfite sequencing (DBS), we screened swim-up sperm samples from 40 normozoospermic and 93 oligoasthenoteratozoospermic (OAT) patients for *H19* and *MEST* methylation. Based on this screening, we defined three patient groups: normal controls (NC), abnormally methylated infertile (AMI; n=7) and normally methylated infertile (NMI; n=86). Whole genome bisulfite sequencing (WGBS) of five NC and five AMI samples revealed abnormal methylation levels of all 50 imprinting control regions in each AMI sample. To investigate whether this finding reflected epigenetic germ line mosaicism or the presence of residual somatic DNA, we made a genome-wide inventory of soma-germ cell specific DNA methylation. We found that >2,000 germ cell-specific genes are promoter-methylated in blood and that AMI samples had abnormal methylation levels at these genes, consistent with the presence of somatic cell DNA. The comparison between the five NC and six NMI samples revealed 19 differentially methylated regions (DMRs), none of which could be validated in an independent cohort of 40 men. Previous studies reported a higher incidence of epimutations at single CpG sites in the CTCF-binding region 6 of *H19* in infertile patients. DBS analysis of this locus, however, revealed an association between DNA methylation levels and genotype (rs2071094), but not fertility phenotype.

**Conclusions:** Our results suggest that somatic DNA contamination and genetic variation confound methylation studies in sperm of infertile men. While we cannot exclude the existence of rare patients with slightly abnormal sperm methylation at non-recurrent CpG sites, the prevalence of aberrant methylation in swim-up purified sperm of infertile men has likely been overestimated, which is reassuring for patients undergoing assisted reproduction.

## Background

Children born following assisted reproductive technology (ART) treatments are thought to have a higher prevalence of imprinting defects [1]. One potential origin of such epimutations may lie in the oocyte and embryo culture, which are commonly part of ART procedures [2]. Apart from that, a number of studies have shown that male infertility itself is associated with aberrant DNA methylation profiles, particularly of imprinted genes [3–6], suggesting that ART may facilitate the transmission of imprinting errors in sperm cells to the next generation. This latter aspect is still, notably, a matter of much debate [7].

Imprinting defects can originate at the different phases of DNA methylation erasure and establishment, occurring during the development of the germline. Sperm originates from primordial germ cells (PGCs). These cells are specified early during embryo development and undergo almost complete erasure of DNA methylation, which allows the establishment of male germline-specific DNA methylation profiles during later stages of gametogenesis [8]. Erasure of DNA methylation in the PGCs takes place in two sequential stages. During the initial stage, a global decrease in methylated cytosines occurs, whereas in the second stage methylation is removed from imprinting control regions (ICRs) and meiotic genes [9]. These phases of methylation erasure result in an epigenetic ground state with methylation levels in PGCs as low as 7-8% at week 11 of human foetal development. The process of *de novo* methylation was found to be re-initiated in PGCs from 19 week-old human foetuses [10]. Primate data suggests that this process continues well after birth in germ cells, which are then termed spermatogonia, and appears to be completed only during puberty [11]. Errors in the process of methylation erasure or re-establishment in a proportion of the PGCs were considered as a possible explanation for subpopulations of sperm displaying aberrant methylation levels in the adult [12, 13]. This explanation is conceivable as those few specified PGCs undergo proliferation and give rise to the population of spermatogonia, which colonize the seminiferous cords of the testes. Apart from the ability to self-renew, spermatogonia can also give rise to differentiating daughter cells through entering spermatogenesis upon puberty. This differentiation process is based on the development of spermatogonial clones, which can result in formation of 16 sperm cells in humans [14, 15]. Incorrect erasure or re-establishment of methylation patterns in individual PGCs could therefore lead to a population of spermatogonia giving rise, via clonal divisions, to a subpopulation of sperm with aberrant methylation profiles.

To address the presence of imprinting errors in sperm, a number of studies have assessed the methylation status of the maternally imprinted gene *MEST* and the paternally imprinted gene *H19* in fertile and infertile men [16]. A meta-analysis suggested a 9.91-fold higher risk ratio for aberrant methylation in the differentially methylated region of *H19* for infertile men. In contrast, no increased risk ratio was found for *MEST* [16].

Careful examination of individual studies suggests four general subgroups of patients, based on the methylation status of *H19* and *MEST*: 1) men with normal *MEST* and *H19* methylation; 2) men with abnormal *MEST* methylation; 3) men with abnormal *H19* methylation; 4) men with impaired methylation patterns in both *MEST* and *H19* [5, 17]. Employing deep bisulfite sequencing at single-allele resolution, a constant proportion of sperm in oligoasthenoteratozoospermic men was found aberrant in four analysed imprinted genes (*H19*, *MEG3* and *MEST*, *KCNQ1OT1*), whereas normozoospermic samples presented as an epigenetically homogenous population [13].

In addition to target gene approaches, individual studies have employed methylation arrays to assess methylation changes at selected CpGs (up to 450,000) that may be present in sperm from infertile men (see for example [18] and [6]). Interestingly, these studies did not report alterations in the imprinted genes *H19* and *MEST* but did identify CpG sites associated with 48 imprinted genes displaying aberrant methylation [6]. Apart from that, a number of additional CpG sites throughout the genome, not associated with ICRs, showed aberrant DNA methylation patterns [6, 18].

As previous studies were largely focused on the analysis of a few imprinted genes and a small fraction of genomic CpG sites, we set out to analyze the genome-wide DNA methylation patterns of human sperm in normozoospermic and oligoasthenoteratozoospermic men. For this, we used a combination of whole genome bisulfite sequencing (WGBS), which provides information on the methylation status of nearly all of the 28,000,000 human CpGs sites, and targeted deep bisulfite sequencing (DBS).

## Results

### Screening of *H19* and *MEST* methylation levels in swim up purified sperm DNA from 93 oligoasthenoteratozoospermic and 40 normozoospermic patients

In order to select patients for whole genome bisulfite sequencing (WGBS) analysis, we measured *H19* (CTCF6 region) and *MEST* methylation levels by deep bisulfite sequencing (DBS) of swim-up purified sperm DNA in a cohort of 40 normozoospermic (Normal) and 93 oligoasthenoteratozoospermic (Infertile) patients (Fig. 1AB, Table 1, Additional file 1: Tables S1 and S2). A principle component analysis (PCA) of *H19* and *MEST* methylation values showed that some infertile men clearly deviated from the remaining samples (Additional file 2: Fig. S1). Since the first principal component (PC1) explains most of the variability of the samples, we considered samples with PC1 score < 0.1 as normally methylated and with PC1 score ≥ 0.1 as abnormally methylated. According to this threshold we subdivided the patients into normally methylated normal controls (NC, n=40), abnormally methylated infertile (AMI, n=7) and normally methylated infertile men (NMI, n=86) (Fig. 1C, Additional file 2: Fig. S1).

**Table 1:** Clinical parameters of the included patient samples

| Clinical parameters | Normal patients<br>(n=40) | Infertile patients<br>(n=93) |
| --- | --- | --- |
| Age (Years) | 35.5 (33-38) | 36 (33-40) |
| Total sperm count (Millions) | 176.6 (107.5-282.4) | 13.1 (6-23) |
| Sperm concentration (Millions/ml) | 39.9 (26.2-69.3) | 3.2 (1.5-5.3) |
| Sperm progressive motility (%) | 50 (44.8-54) | 36 (26-46) |
| Sperm normal forms (%) | 5 (4-6) | 2 (1-3) |
| Vitality (%) | 70 (66-79) | 61 (52.8-67.2) |
| FSH (U/l) | 3.2 (2.5-3.9) | 6.4 (4.2-9.5) |
| LH (U/l) | 2.8 (2.2-3.5) | 3.7 (2.6-5.1) |
| Testosterone (nmol/l) | 18.1 (14.9-21.6) | 16.6 (12.4-22.8) |

Median values and the 25th and 75th percentile are given for the different parameters.

**Fig. 1:**
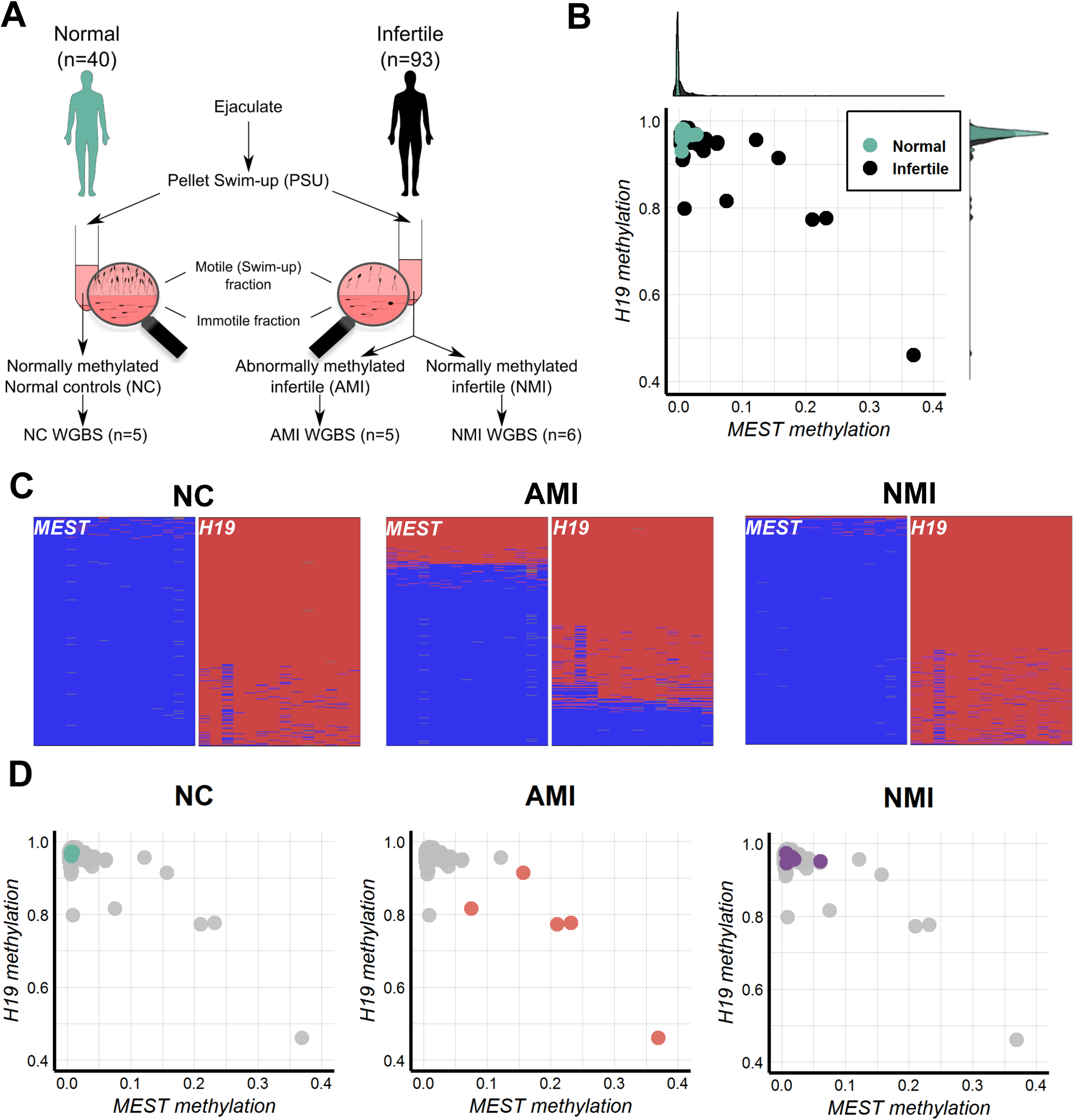
Sperm samples selection for WGBS and establishment of groups based on *MEST* and *H19* methylation. A) Schematic representation of the experimental design. B) Dotplot representing the mean methylation levels of *MEST* and *H19* measured by deep bisulfite sequencing in 40 normal (teal) and 93 infertile (black) sperm samples. At the margins, two density plots show the distribution of the *MEST* and *H19* mean methylation values in the normal and infertile cohort of samples (Additional file 1: Table S2). C) Example of deep bisulfite sequencing results of *MEST* and *H19* in the three groups: normal control (NC), abnormally methylated infertile (AMI) and normally methylated infertile (NMI). Each horizontal line of a plot represents a unique sequence read, while each vertical position represents a CpG site (methylated sites in red, unmethylated sites in blue). D) Mean methylation values for *MEST* and *H19* in the five NC (teal), five AMI (orange) and six NMI (purple) selected for the WGBS.

### Whole methylome analysis of swim-up sperm DNA from patients with normal and impaired spermatogenesis

For WGBS we chose five NC, five AMI and six NMI samples (Fig. 1D). Following the recommendations by Ziller et al. [19], we sequenced the samples at 13-16× coverage (Additional file 1: Table S3). For comparative analyses, we used previously generated WGBS data of isogenic blood and sperm samples of 12 normozoospermic men (two pools of six individuals each) [20].

### Evaluation of whole methylome data for the 50 known imprinting control regions

In order to determine whether, in addition to *MEST* and *H19,* other imprinted loci were also affected by aberrant methylation, we analysed the WGBS methylation values for the 50 known maternally and paternally methylated ICRs [21]. We found that the five AMI samples had abnormal methylation levels at all ICRs and that the degree of aberrant methylation at these regions was highly correlated within each sample (Fig. 2, Additional file 2: Fig. S2 and Additional file 1: Table S4). In contrast, the ICR methylation values for the six NMI samples were similar to the observed values in NC samples. Moreover, a PCA of the 15 methylomes revealed that the AMI samples span across the PC1 axis, while NC and NMI samples group together and in the opposite extreme compared to the blood samples (Additional file 2: Fig. S3).

**Fig. 2:**
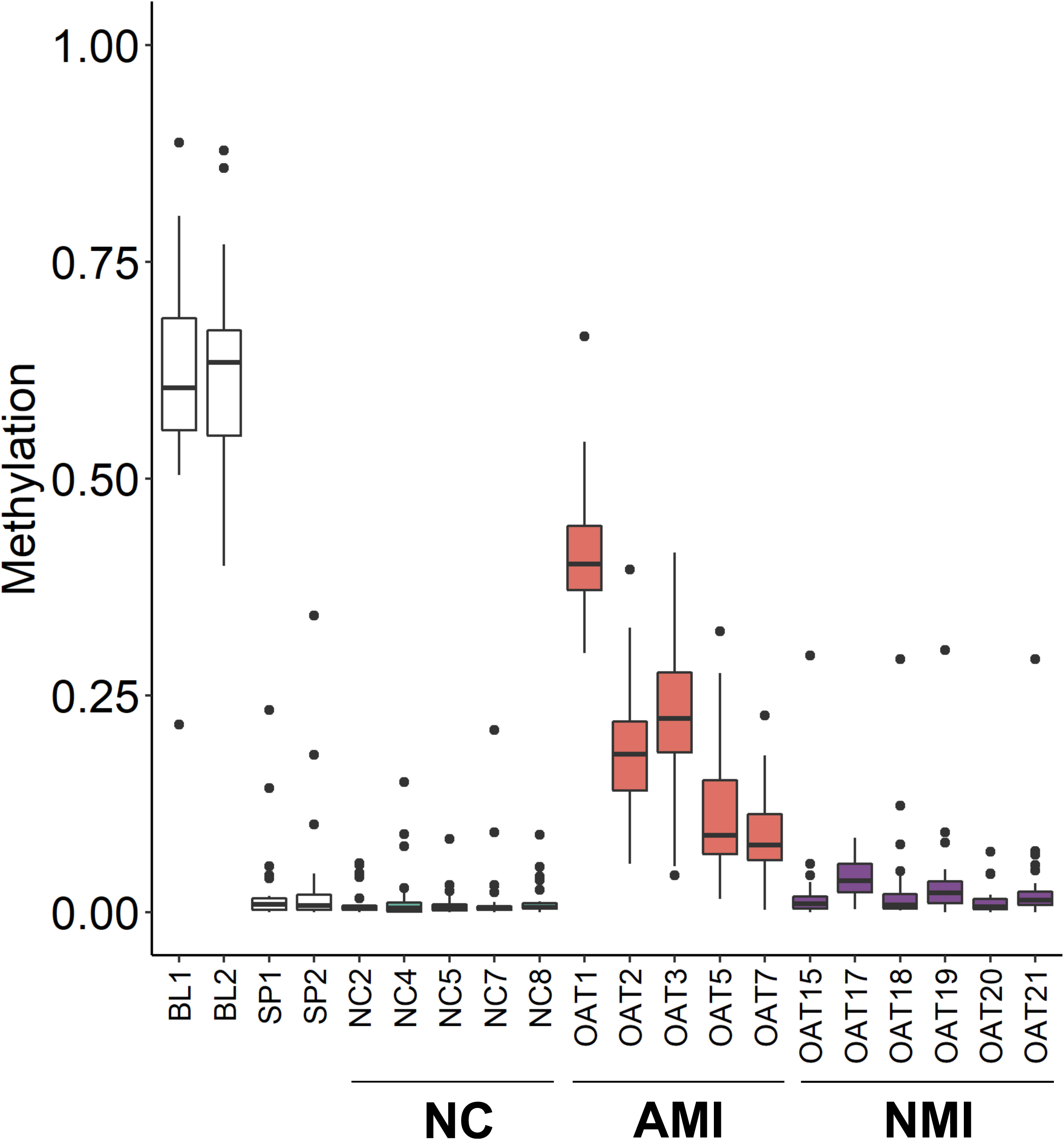
Methylation levels of the oocyte genomic imprints. Box plots showing the distribution of 34 oocyte DMRs methylation values in blood and sperm DNA (Additional file 1: Table S4). Datasets from Laurentino et al. [20] appear in white (BL1 and BL2 – blood, SP1 and SP2 – sperm), NC sperm samples in teal, AMI sperm samples in orange and NMI sperm in purple. Box plots elements are defined as follows: center line: median; box limits: upper and lower quartiles; whiskers: 1.5× interquartile range; points: outliers.

### Inventory of differentially methylated regions between sperm and blood derived somatic cells

To investigate whether the aberrant methylation levels in the AMI group reflect epigenetic germ line mosaicism or the presence of previously undetected somatic DNA, we made an inventory of soma-germ cell specific methylation differences. For this, we compared published WGBS data of isogenic blood and sperm samples of 12 normozoospermic men [20] with two different bioinformatic tools (camel and metilene) to identify methylation differences. By defining a differentially methylated region (DMR) as a region of at least 10 CpGs with a methylation difference of at least 80% and a minimum coverage of five reads, we detected 32,686 DMRs, of which 6,159 overlap the promoter of 5,892 genes (Fig. 3A). Of these genes, 2,462 were among the 8,175 genes previously shown to be expressed in germ cells and not in testicular somatic cells [22] and which are putatively regulated by DNA methylation of 2,764 DMRs (Additional file 1: Table S5). In line with the expression analysis, almost all of these gene promoters were methylated in blood and unmethylated in sperm. Analysis of the methylation levels of the 2,764 DMRs revealed that the five AMI samples have aberrant methylation at all soma-sperm specific differentially methylated genes (Fig. 3B, Additional file 2: Fig. S4 and Additional file 1: Table S5). Moreover, in each sample, the degree of aberrant methylation was similar to the levels observed for the imprinted regions (Fig. 2, Additional file 2: Fig. S2 and Additional file 1: Table S4).

**Fig. 3:**
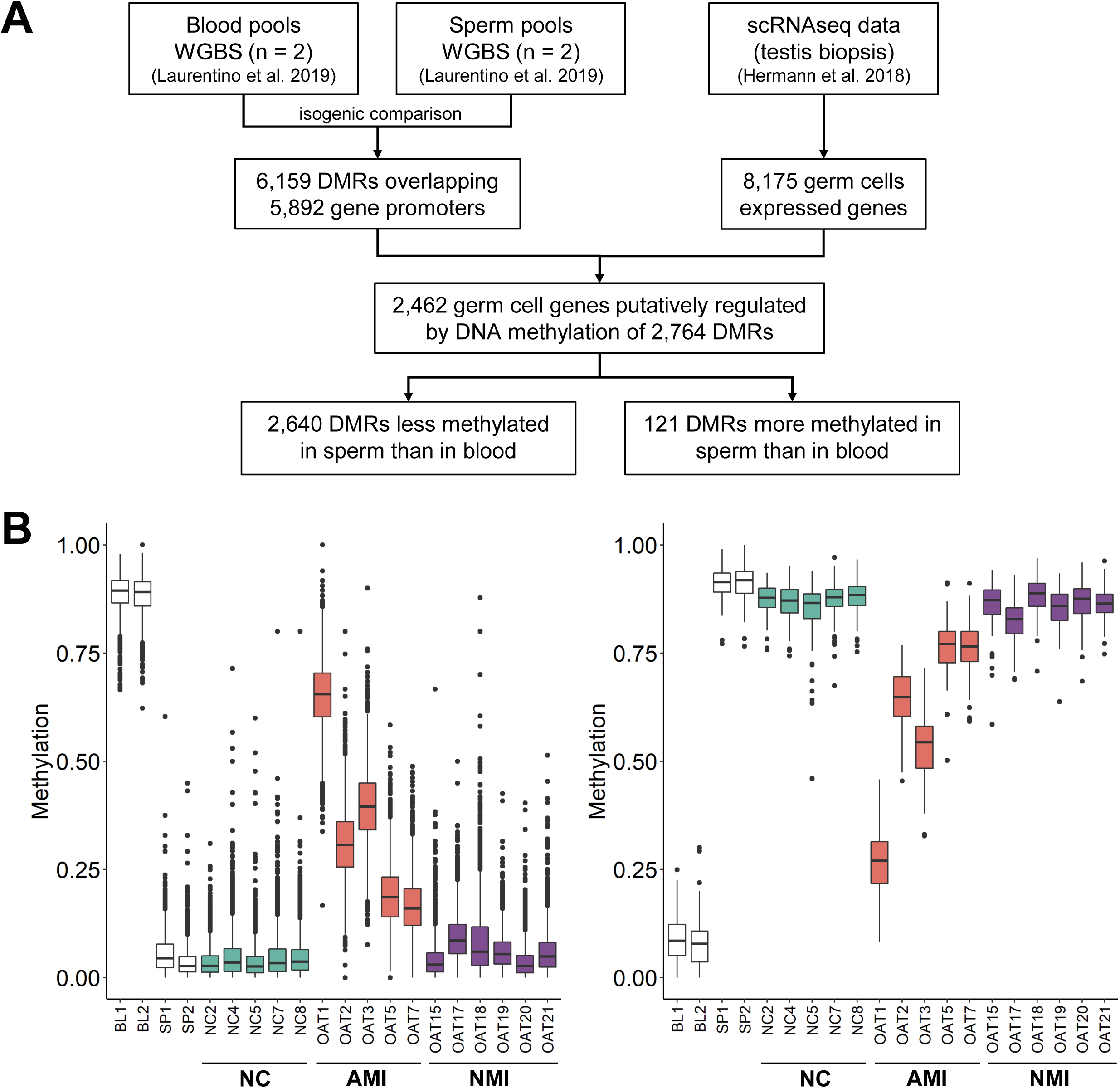
Inventory of the sperm-soma DMRs putatively regulating promoters of 2,462 testicular germ cell-specific genes. A) Flow chart of the discovery of 2,764 sperm-soma DMRs using blood datasets as somatic representatives. B) Box plots showing the distribution of the methylation values of 2,640 DMRs less methylated in sperm than in blood (left) and 121 DMRs more methylated in sperm than in blood (right) (Additional file 1: Table S5). Datasets from Laurentino et al. [20] appear in white (BL1 and BL2 – blood, SP1 and SP2 – sperm), NC sperm samples in teal, AMI sperm samples in orange and NMI sperm samples in purple. Box plots elements are defined as follows: center line: median; box limits: upper and lower quartiles; whiskers: 1.5× interquartile range; points: outliers.

Most recently, Luján et al. claimed to have identified 217 DMRs useful for fertility assessment. In their study, they analysed unpurified sperm samples by methylation-dependent immunoprecipitation (MeDIP-Seq) [23]. We determined the methylation levels of these DMRs in the blood-sperm WGBS dataset [20] and our five NC and six NMI samples (Additional file 1: Table S6). We found that the DMRs are unable to distinguish sperm of fertile men from sperm of infertile men (Additional file 2: Fig. S5). Rather, they discriminate between clean sperm samples and sperm samples containing somatic DNA, as the 50 ICR DMRs and our inventory of 2,764 soma-sperm DMRs do, but the latter do so with higher sensitivity (Fig. 2 and Fig. 3).

To further validate the findings in our patients, we performed DBS for *XIST* and *DDX4* loci, previously shown to be fully unmethylated in normal sperm [24], on the 40 normal controls and the 93 infertile patient samples used in the initial screening (Additional file 1: Table S2). We further confirmed that each of the five AMI that were subjected to WGBS showed an aberrant methylation level at these two loci, which was highly correlated with the aberrant methylation in both the imprinted regions and the soma-sperm specific differentially methylated genes. All normal controls and 77 of the normally methylated infertile were found to have the expected *XIST* and *DDX4* methylation levels (<6%; Additional file 2: Fig. S6). From the two AMI samples not analysed by WGBS, one (SOAT7) was shown to have *DDX4* methylation levels consistent with the presence of somatic cell DNA (Additional file 1: Table S2). The other (SOAT6) showed aberrant methylation levels for *H19* CTCF6, but was considered normal for *MEST*, *XIST* and *DDX4* (Additional file 2: Fig. S7). This sample had a similar pattern in the CTCF4 region of *H19*, but the fraction of completely unmethylated reads was smaller. We sequenced additional ICRs and compared the DBS methylation levels in this sample (Additional file 2: Fig. S7) with that of a representative NC (VN25, Additional file 2: Fig. S8). SOAT6 has a very small proportion of completely methylated reads in the *XIST*, *KCNQOT1* and *PEG10* amplicons. In summary, we conclude that despite swim-up purification, somatic cell DNA was still present in some NMI and AMI samples and therefore these samples were excluded from further analysis.

### Identification of differentially methylated regions in sperm from normal and oligoasthenoteratozoospermic men

To identify true DMRs between the sperm of fertile and infertile men, we compared the genome-wide methylomes of six NMI and five NC sperm samples that are devoid of somatic DNA. (Fig. 4A). Using two different bioinformatic tools, we identified 103 DMRs with at least five CpGs, a methylation difference of at least 0.3 and a minimum coverage of five reads (Additional file 1: Table S7). Since the genetic background (i.e. DNA polymorphisms) may affect DNA methylation [25], some DMRs may display a higher range of values within a group. Therefore, to reduce the potential influence of the genetic background, we limited the range of methylation values within the normozoospermic group to 0.3, thus keeping 19 of the 103 DMRs (Fig. 4A and Additional file 1: Table S7). Three of the 19 DMRs were hypermethylated in normozoospermic samples, while the remaining 16 were hypermethylated in the NMI patients (Fig. 4B and Additional file 1: Table S7).

**Fig. 4:**
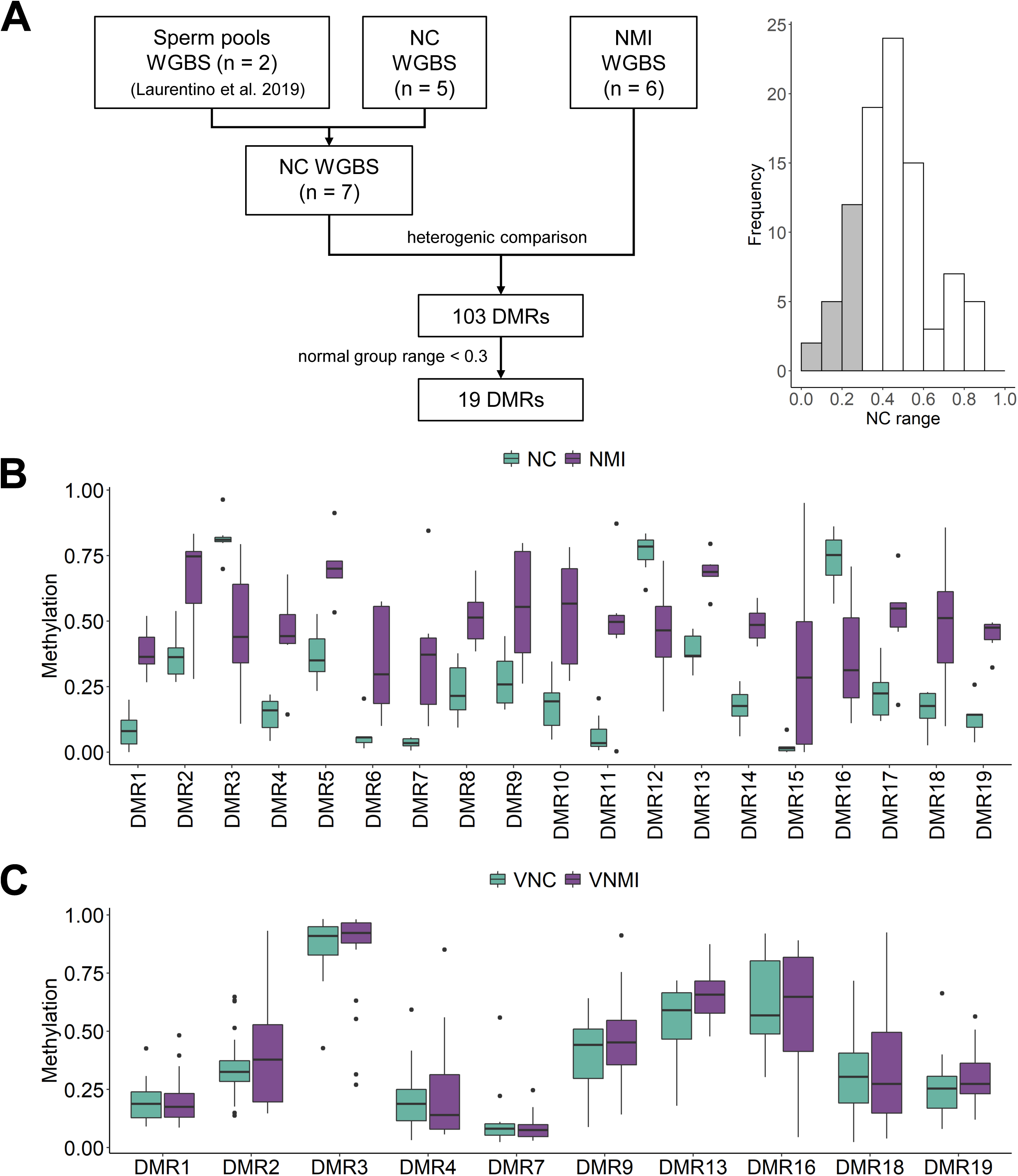
Discovery of normal controls vs. normally methylated infertile sperm DMRs. A) Flow chart of the discovery of 19 DMRs between NC and NMI sperm groups (left) and the distribution of DMRs according to the range of values in normal controls, with DMRs having NC range < 0.3 highlighted in grey (right). B) Box plots showing for the 19 DMRs the distribution of the methylation values for NC (teal, n = 5), and NMI (purple, n = 6) (Additional file 1: Table S7). C) Box plots showing for 17 DMRs the distribution of the methylation values obtained by targeted DBS in an independent cohort. VNC, validation normal control samples (teal, n = 20), VNMI, validation normally methylated infertile samples (purple, n = 20) (Additional file 1: Table S11). Statistical analyses showed no differences between the two groups (Additional file 1: Table S12). Box plots elements are defined as follows: center line: median; box limits: upper and lower quartiles; whiskers: 1.5× interquartile range; points: outliers.

In order to validate the DMRs in an independent cohort, we established reliable targeted DBS assays for 17 DMRs (Additional file 1: Table S8; specific primers could not be designed for DMR6 and DMR12 due to the presence of highly homologous sequences in the genome). Although the DBS approach targets only DMR CpG subsets (coordinates in Additional file 1: Tables S7 and S8), the distributions of WGBS and WGBS CpG subset methylation values are the same (Additional file 1: Tables S9 and S10). Due to the limited amount of infertile sperm DNA, we first analysed 20 normal control samples (VNC) and then selected DMRs for further validation in 20 normally methylated infertile swim-up sperm DNA samples (VNMI). After sequencing the VNC samples for each of the 17 DMRs, we selected 10 DMRs based on the number of VNMI methylation values outside of the normal samples methylation range (Additional file 2: Fig. S9 and Additional file 1: Table S11). Following sequencing each of the 10 selected DMRs in the 20 VNMI samples and comparison with the VNC data, none of the DMRs could be validated (Fig. 4C and Additional file 1: Tables S11 and S12).

### Influence of single nucleotide polymorphism (SNPs) on *H19* methylation levels

Single CpG sites in the CTCF6 binding site of *H19* have previously been shown to be differentially methylated in normal and infertile patients [4, 17, 26–32]. In order to analyse this further, we performed a PCA, using as loadings the methylation values of the individual 14 CpG sites analysed by DBS in all the individuals showing no presence of somatic cell DNA according to *XIST* and *DDX4* assay results (n=118, NC=40, NMI=77, AMI=1; Additional file 1: Table S13). This analysis showed that the variation in PC1 was mainly due to the CpG3 methylation levels (Additional file 2: Fig. S10). The peculiarity of CpG3 is also visible in the amplikyzer plots (Fig. 1C, Additional file 2: Fig. S7 and S8). CpG3 is in the vicinity of a G/A-SNP (rs10732516; Fig. 5A). Since the genotype of this SNP is masked by bisulfite treatment, we used the nearby rs2071094 SNP, which is in high linkage disequilibrium (r^2^=0.99 and D’=1 according to annotations by HaploReg v4.1 [33]) to investigate the possible effects of these SNPs on *H19* methylation values. Such an effect has previously been reported in blood and placenta [34, 35]. We observed that individuals clustered in the PCA according to their rs2071094 genotype (AA, AT, TT) (Fig. 5B). TT men showed a significantly lower CpG3 methylation compared to the individuals with AT or AA genotype (Fig. 5C), and AT men showed a significantly lower methylation in the reads containing the T allele compared to the A (Fig. 5D and Additional file 1: Tables S14 and S15). Finally, the subdivision of patients according to the diagnosis (NC or NMI) did not show any significant difference between normal and infertile patients sharing the same genotype (Fig. 5E). This shows that the methylation level of CpG3 is affected by genetic variation irrespective of the fertility status.

**Fig. 5:**
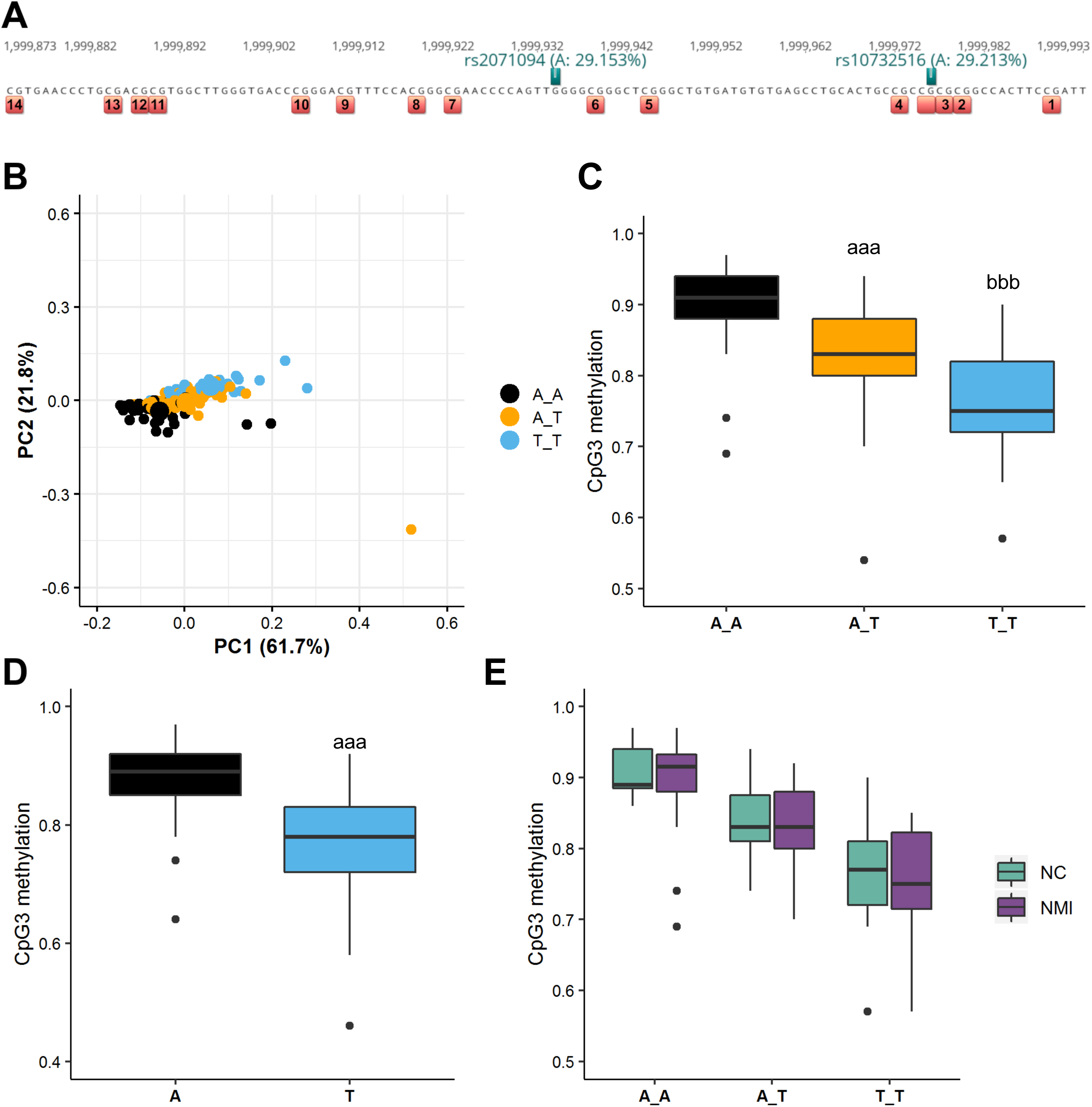
Analysis of the influence of the SNP rs2071094 on the *H19* methylation levels. A) Schematic representation of the *H19* CTCF6 locus showing the CpGs analyzed by DBS (red and numbers 1-14) and the SNP-masked CpG (red). SNPs in high linkage disequilibrium are shown in green. Numbers on top refer to hg38 coordinates of chromosome 11. B) Principal component analysis (PCA) of the 14 CpG sites in the *H19* locus obtained by DBS for the 40 normal controls (NC), 77 normally methylated infertile (NMI) and one abnormally methylated infertile (AMI) color-coded according to the SNP rs2071094 genotype: A/A black, A/T orange, T/T lightblue. C) Box plot showing the distribution of the CpG3 methylation in the 118 patients subdivided according to the SNP rs2071094 genotype. Statistically significant differences are denoted by letters: a—AT different from AA, b—TT different from AT and AA. P-values are denoted by the number of letters, e.g. aaa p < 0.001 (Additional file 1: Table S15). D) Box plot showing the CpG3 methylation in the A versus the T allele of the 49 AT patients (Additional file 1: Table S14). E) Box plot showing the CpG3 methylation in the 40 normal controls (NC, teal) and 77 normally methylated infertile (NMI, purple) divided according to the SNP rs2071094 genotype.

## Discussion

Aberrant DNA methylation patterns of imprinted genes have been reported in semen samples from infertile men in a number of studies [16, 36]. While the majority of studies focused on the analysis of selected ICRs, mainly *MEST* and *H19,* these reports still differed with regard to the observed differences between normal and infertile men. Specifically, aberrant methylation patterns for only *MEST* or *H19* were described in some patients, whereas others apparently carried a subpopulation of sperm which showed the same degree of aberrant imprinting in multiple imprinted genes (*MEST*, *LIT1, H19*, *MEG3)* and thereby indicated epigenetic mosaicism in sperm from OAT men [13]. As these epigenetic aberrations might be transmitted to the offspring, it was a clinical necessity to assess the extent of these aberrations, not only for selected genes, but also for the entire genome. To this end, this study sought to assess the DNA methylation levels in normal and severely impaired spermatogenesis by whole genome and ultra-deep bisulfite sequencing.

In the screening process of the 93 samples from patients with severe oligoasthenoteratozoospermia, which make our study one of the largest in its field, only 1% showed aberrant methylation for *MEST* and also 1% for *H19*. Five per cent of samples appeared to be aberrantly methylated at both imprinted genes, whereas the great majority (93%) showed normal methylation levels for *MEST* and *H19*. The presence of these four subgroups and the distribution among them when analysing only *MEST* and *H19* methylation values is in line with previous publications [5, 17], although percentages of samples with aberrant profiles were generally higher (e.g. 57% in Poplinski et al. [5]).

While in some studies, all of the analysed CpG sites within the CTCF6 region of *H19* were either methylated or unmethylated [5], in other studies the methylation differences were restricted to single CpG sites within this region [4, 17, 26, 27, 29, 31, 32]. We also observed, in both normozoospermic and OAT samples, a fraction of partially unmethylated reads in our *H19* DBS amplicons, which cover a large proportion of the CpGs that had been analysed by Sanger sequencing of subcloned PCR products or by pyrosequencing in the above mentioned studies. The most variable *H19* CpG in our assay is CpG3, which corresponds to CpG4 in Camprubi et al. [27], CpG5 in Boissonnas et al. [26], and CpG6 in other studies [4, 17, 29, 31, 32]. We demonstrate that variation in DNA methylation at this CpG site is correlated with the genotype of a nearby SNP (rs2071094), irrespective of the fertility status, with TT homozygotes having the lowest methylation level and AA homozygotes the highest methylation levels. rs2071094 is in high linkage disequilibrium with CpG-SNP rs10732516 suggesting that the presence or absence of an additional CpG site next to CpG3 could influence the methylation of the latter. These results support the view that DNA methylation patterns are influenced to a large extent by the genetic background [25] and suggest that studies reporting reduced methylation levels of this CpG in infertile men might have been confounded by a fortuitously higher T allele frequency in cases compared to controls. We identified one individual sample showing an aberrant methylation level of the *H19* CTCF6 and CTCF4 regions as well as a very small proportion of completely methylated reads in the *XIST*, *KCNQOT1* and *PEG10* amplicons. We are uncertain whether this sample carries a true *H19* epimutation, has a rare genetic variant or contains minute amounts of somatic DNA, which show up in some but not in all PCRs.

In this study, we focused on the genome-wide DNA methylation analysis of the two most prominent groups of infertile samples: those, with abnormal methylation of *MEST* and *H19* (AMI), and those with normal methylation levels in both regions (NMI). Unexpectedly, the former group of samples displayed the same level of aberrant methylation not only in *H19* and *MEST*, but in all of the 50 known ICRs as well as in *DDX4* and *XIST*. Moreover, 2,764 soma-germ cell specific DMRs were also aberrantly methylated to the same degree. A scenario of such comprehensive reprogramming failure appears highly unlikely. In contrast, the presence of residual somatic cell DNA, shifting the methylation level towards that of somatic cells, appears to be the more plausible explanation. After exclusion of samples showing a clear presence of somatic DNA (16% of our infertile samples), only one sample with aberrant methylation at *H19* remained. Although it is unclear whether this sample contains traces of somatic DNA, the percentage of infertile patients possibly carrying an imprinting defect in our cohort (0-1%) is much lower than previously reported (as high as 57% in Poplinski et al. [5]). We suspect that other studies also suffer from DNA contamination issues.

The origin of somatic cell DNA in swim-up purified sperm samples remains hitherto unclear. It has been reported that increased numbers of leucocytes are present in the semen of 30% of infertile men, even in the absence of an infection [37]. It appears possible that in these cases, somatic cells or cell fragments that escape quality controls could be amidst the very few sperm that are present in the infertile samples and skew the analyses in the direction of a somatic cell profile. This unexpected result highlights the importance of assessing sperm DNA samples for absence of somatic cell DNA prior to methylation studies. Along this line, pre-screening approaches have been published, which describe multiple sites enabling the distinction of germ cell versus somatic cell derived DNA [38]. Here, we describe 2,764 DMRs that overlap with the promoters of 2,462 genes previously shown to be expressed in germ cells and not in testicular somatic cells [22]. This comprehensive list of DMRs constitutes a valuable resource for future studies seeking to assess the purity of their sperm samples.

It is surprising that so many genes, both protein and non-protein coding genes, appear to be regulated by promoter methylation. Most often, cellular differentiation does not involve promoter methylation, but methylation of distal regulatory elements such as enhancers. Interestingly, most of the 2,462 genes are methylated in blood cells and unmethylated in germ cells. This suggests that these genes need to be permanently silenced in somatic cells. Since many of these genes play a role in meiosis, it is tempting to speculate that these genes are permanently silenced in somatic cells to prevent them from interfering with mitosis.

When comparing the genome-wide methylomes of sperm samples from fertile and infertile patients displaying normal *MEST* and *H19* methylation levels we did not find any recurrent methylation difference between the two groups. This is in contrast to a recent report in which the authors claim to have identified 217 DMRs between unpurified sperm from nine fertile and 12 infertile men [23]. However, as shown here, the methylation levels at these regions reflect the admixture of somatic DNA and are not biomarkers of infertility.

Our findings show that the DNA methylation patterns of clean sperm are normal, which is reassuring for patients undergoing ART treatment. It is possible that spermatogonia with DNA methylation abnormalities exist, but they likely do not contribute to the mature, swimming sperm population, suggesting that cells with an abnormal methylome may be counter-selected at later stages of spermatogenesis. It is of note though, that we only considered regions consisting of more than five CpG sites for our analysis, which is in contrast to previous publications performing array analysis and considering individual CpG sites [6, 18]. It should be noted, however, that aberrant methylation restricted to one or a few CpGs of an ICR, if real, is unlikely to be of clinical relevance, because in all patients with an imprinting disease based on imprinting errors, almost all CpGs of an ICR are affected [39, 40].

## Conclusions

Our results suggest that the undetected presence of somatic DNA as well as genetic variation confound methylation studies in sperm of infertile men. After controlling for these confounders, we have found no evidence for recurrent epimutation in imprinted genes or elsewhere in the genome in sperm of severely oligoasthenoteratozoospermic men. While we are aware that WGBS is underpowered to detect rare patients with slightly abnormal sperm methylation levels at non-recurrent CpG sites, we conclude that the prevalence of aberrant methylation in infertile men has likely been overestimated, which is reassuring for patients undergoing ART treatment. In the course of this study, we have also found that a large number of germ cell-specific genes are regulated by promoter methylation. The list of soma-germ cell specific DMRs can be used for assessing the quality of sperm preparations and for studying the epigenetic regulation of spermatogenesis in more details.

## Methods

### Sample selection and clinical information

The patients included in this study were selected among those attending the Department of Clinical and Surgical Andrology at the Centre of Reproductive Medicine and Andrology (CeRA, Münster, Germany) for fertility treatment. All the patients underwent full physical evaluation and those with known genetic causes of infertility, chromosomal aberrations, under pharmacological treatment, with a history of cryptorchidism, acute infections and tumours were excluded from the analysis. Blood samples were taken for hormone measurements including gonadotropins and testosterone following published protocols [24]. Moreover, semen analysis was performed according to the WHO manual [41]. In total, 133 individuals were selected and subdivided into two age matched groups according to the spermiogram results: 40 normal controls (NC) diagnosed as normozoospermic and 93 diagnosed as oligoasthenoteratozoospermic, oligoteratozoospermic or oligozoospermic, which are termed OATs throughout the manuscript (Additional file 1: Table S1).

### Swim-up procedure for isolation of motile sperm

Swim-up procedure was used to isolate the motile sperm cells, in line with preparation of samples for assisted reproductive technology treatment. Briefly, after an incubation period of 30 minutes (min) at 37°C, 1-2 ml of ejaculate were mixed with the same amount of sperm preparation medium (Origio, Denmark), by using a cell culture tested disposable pipette. The mixture was then centrifuged at 390 g for 10 min, the supernatant decanted and the remaining drops aspirated. The pellet was washed with 2 ml of medium and centrifuged at 390 g for 10 min. After removing the supernatant 1 ml of medium was carefully added to the pellet in order to not dissolve or wash it off. As a precaution the tube was briefly centrifuged for 1 minute at 390 g and then incubated for 60 min at 37°C and 5% CO_2._ After 1 hour of incubation, 500-700 µl of the uppermost layer were collected and stored in a small cell culture tube. 20 µl of the cell suspension were used to determine the sperm concentration in a Neubauer improved counting chamber (Additional file 1: Table S1).The rest of the volume was further centrifuged for 5 min at 16,060 g, the supernatant was discarded and the sperm pellet was stored at −20°C.

### DNA isolation and bisulfite conversion

The DNA isolation was performed on the swim up purified sperm using the MasterPure DNA purification kit (Epicentre Biotechnologies, Madison, WI, USA) as previously described [13]. DNA concentration was measured using a fluorescence plate reader (FLUOstar Omega, BMG Labtech, Germany). The bisulfite conversion was performed on 100 ng of sperm DNA using the EZ DNA Methylation-Gold kit (Zymo Research, Freiburg, Germany) according to the manufacturer’s protocol. The bisulfite converted DNA was eluted in 10 µl of TE buffer.

### Whole genome bisulfite sequencing

Sperm WGBS libraries were prepared according to the tagmentation-based method described by Souren et al. [42] with some modifications. Briefly, 10 ng sperm DNA supplemented with 1% unmethylated lambda-DNA (Promega) were incubated in a 50 µl reaction with 0.8 µl of Tn5 transposase at 1× TD buffer from the Nextera library preparation kit (Illumina) for 5 min at 55°C. Tagged DNA was purified with the DNA Clean & Concentrator-5 kit (Zymo Research) eluting with 14 µl EB buffer (Qiagen), followed by gap repair by adding 2 µl of 10× NEBuffer 2 (NEB), 3 µl of dNTPs (2.5 mM each) and 5 U Klenow exo- (NEB) and incubating for 1 h at 30°C. Bisulfite conversion was performed using the EZ DNA Methylation-Gold kit (Zymo Research) according to the manufacturer’s instructions. Indexed-libraries were obtained by enrichment PCR with 1x HotStarTaq Master Mix (Qiagen), 100 nM of each primer and 10 µl bisulfite-converted DNA in 40 µl reactions (PCR settings: 95°C 15 min, 12× (95°C 30 s, 53°C 2 min, 72°C 1 min), and 72°C 7 min). Reactions were purified twice using 0.8x volume AMPure XP Beads (Beckman Coulter) and eluted in 10 µl EB buffer (Qiagen). Libraries were sequenced in HiSeq4000 100-bp paired-end runs (Illumina) using one lane per sample.

### WGBS data analysis

Raw read data was aligned against reference genome hg38 using bwa-meth [43] (v0.2.0) and deduplicated by Picard [44] (v2.18.15). We used MethylDackel [45] (v0.3.0) for subsequent methylation calling. For quality control we used MultiQC [46] to integrate quality metrics collected by Picard, FastQC [47] (v0.11.8) and Qualimap [48] (v2.2.2b). We chose camel [49] (v0.4.7) and metilene [50] (v.0.2.6) to call DMRs. Average coverage per DMR was computed using mosdepth [51] (v0.2.3). We filtered DMRs based on the number of CpGs covered, methylation differences between groups, q-values reported by metilene and average coverage. For the blood/sperm comparison we required DMRs to cover at least 10 CpGs with at least 80% difference in methylation, minimum coverage of 5 reads and a maximum q-value of 0.05. When comparing NC and OAT samples we set the thresholds to 5 CpGs, 30% methylation difference, minimum coverage of 5 and a maximum q-value of 0.05. After filtering we merged DMRs using the GenomicRanges R package [52]. Merged DMRs were annotated for overlap with CGIs using data from the UCSC database [53]. Genes and promoters were annotated using information from the Ensembl database [54]. We require genes to be marked as either protein coding, long non-coding RNA or miRNA. Promoters were defined as the 2000 bp region around TSSs.

### Targeted deep bisulfite sequencing

Targeted DBS was performed on the Roche/454 or the Illumina MiSeq platform essentially as described previously [55] using 100 ng of sperm DNA for bisulfite conversion, the primer pairs and PCR conditions described in the Additional file 1: Table S8. For the *H19* amplicon, although it comprises 15 CpGs, only 14 CpGs are shown since the CpG affected by a known polymorphism (rs10732516) was masked in the analyses.

### Statistics

Normality and homoscedasticity tests were performed for all variables and difference between groups was assessed by non-parametric tests: Wilcoxon signed rank test for two dependent groups and Mann-Whitney U test for two independent groups, followed by Bonferroni correction for multiple testing. Kruskal-Wallis rank sum test was used to compare three or more independent groups, followed by multiple pairwise-comparisons. Statistical analysis and graphs plotting were performed using R 3.5.3 [56] and appropriate R packages.

## List of abbreviations

AMI: abnormally methylated infertile
ART: assisted reproductive technology
DBS: deep bisulfite sequencing
DMR: differentially methylated regions
ICR: imprinting control region
NC: normal controls
NMI: normally methylated infertile
OAT: oligoasthenoteratozoospermic
PCA: principle component analysis
PGC: primordial germ cell
SNP: single nucleotide polymorphism
WGBS: whole genome bisulfite sequencing

## Declarations

### Ethics approval and consent to participate

Samples were included in this study following the approval of the Ethics Committee of the University and the Westphalian medical board (reference number of Institutional Review Board: 4INie) and written informed consent.

### Consent for publication

Not applicable.

### Availability of data and materials

The datasets generated in the current study are available in the European Nucleotide Archive (ENA) under the accession number PRJEB34432 and the following link: https://www.ebi.ac.uk/ena/browser/view/PRJEB34432

## Authors’ contributions

NN, SL and BH designed the study; SK supervised clinical patient care and documentation in the database Androbase; SK was responsible for clinical and laboratory testing and data transfer including semen sample preparations for the study; SDP and SL selected the samples based on clinical diagnosis; SDP performed the DNA isolation, bisulfite conversion and screening using deep bisulfite sequencing; SDP, SL and EL analysed the samples by deep bisulfite sequencing; EL prepared the WGBS libraries; MW and MD processed the WGBS data; EL and BH analysed the WGBS data; SDP and EL performed the statistical analyses; EL, SDP, SL, SK, NN and BH interpreted the results and wrote essential parts of the manuscript. All authors approved the final version of the manuscript.

## Acknowledgements

We thank Elisabeth Lahrmann as well as Raphele Kürten, Daniela Hanke, Jolanta Körber, Elena Plester, Sabine Strüwing and Sabine Forsthoff, from the andrology laboratory of the CeRA, for the excellent technical support. We thank Claudia Krallmann, MD (CeRA) for the excellent clinical support. We thank Dr. Janine Altmüller and Christian Becker (Cologne), for sequencing the WGBS libraries, Dr. Christopher. Schröder, Essen, for help in establishing the WGBS data analysis pipeline, Dr. Ludger Klein-Hitpass, Essen, and Dr. Jochen Seggewiß, Carolin and Christian Ruckert (Münster) for sequencing the DBS libraries, and Prof. Dr. Jörg Gromoll, Prof. Dr. Frank Tüttelmann and Prof. Dr. Stefan Schlatt (Münster) for helpful discussions.

## Additional files

Additional file 1: **Table S1:** Clinical parameters. **Table S2:** *MEST*, *H19*, *XIST* and *DDX4* methylation values obtained by DBS for the 133 sperm samples. **Table S3:** WGBS statistics. **Table S4:** WGBS methylation values for the 50 imprinting controls regions. **Table S5:** Annotations and WGBS methylation values for the 2764 soma-sperm DMRs. **Table S6:** WGBS methylation values for the Luján et al. 2019 DMRs. **Table S7:** Annotations and WGBS methylation values for the NC-OAT DMRs. **Table S8:** Primers for generating amplicons for targeted bisulfite sequencing. **Table S9:** Average WGBS methylation values (minimum coverage 5) for the subset of CpGs analysed by DBS in 17 NC-NMI DMR. **Table S10:** Statistical comparison of WGBS and WGBS CpG subset values (WGBS_s5). **Table S11:** Average DBS methylation values in 17 NC-NMI DMR. **Table S12:** Statistical tests concerning NC-NMI DMRs. **Table S13:** Methylation values of the 14 CpG sites in the *H19* locus. **Table S14:** CpG3 methylation values in the A and T alleles in the AT group. **Table S15:** Statistical tests concerning individual CpG sites of *H19*. (XLSX 1,102 kb)

This file is also available at: https://mfga.uni-muenster.de/publication/details?publicationId=5957038

User name: spermEpigenomeInPatientsWithSeverelyImpairedSpermatogenesis

Password: rs2071094

Additional file 2: **Fig. S1:** Principle component analysis (PCA) of *MEST* and *H19* methylation values obtained by DBS for the 133 sperm samples (Additional file 1: Table S2). Samples with PC1 score < 0.1 were considered normally methylated and with PC1 score ≥ 0.1, abnormally methylated. While normal controls (NC) are a homogeneous group of normally methylated samples, infertile sperm samples were subdivided in two groups according to this PC1 threshold (AMI, abnormally methylated infertile; NMI, normally methylated infertile). **Fig. S2:** Methylation levels of the 50 imprinting control regions. Line diagrams showing comparisons between blood (BL1, BL2) and sperm (SP1, SP2) datasets from Laurentino et al. [20] (upper panel), between NC and AMI sperm (middle) and NC and AMI sperm (lower panel) (Additional file 1: Table S4). * Not imprinted according to this data, ** Possible polymorphism. **Fig. S3:** PCA generated for ∼8.7 million CpG loci where all samples show methylation values. Only loci with minimum coverage of five in all samples and minimum mapping quality of 10 are considered. Datasets from Laurentino et al. [20] are shown in white (BL1 and BL2 – blood, SP1 and SP2 – sperm), NC sperm samples in teal, AMI sperm samples in orange and NMI sperm in purple. **Fig. S4:** Methylation levels of 2,761 sperm-soma DMRs. Line diagrams showing comparisons between blood (BL1, BL2) and sperm (SP1, SP2) datasets from Laurentino et al. [20] (upper panel), between NC and AMI sperm (middle) and NC and NMI sperm (lower panel). The 2,640 DMRs less methylated in sperm than in blood are towards the left and the 121 DMRs more methylated in sperm than in blood are on the right (Additional file 1: Table S5). Fig. S5: Methylation levels of the 217 DMRs claimed by Luján et al. [23] to be useful for infertility assessment. Box plots showing the distribution of methylation values for the DMRs stated to be hyper- (190 DMRs, left) or hypomethylated in sperm from infertile vs. fertile men (20 DMRs, right) (Additional file 1: **Table S6**). Datasets from Laurentino et al. [20] are shown in white (BL1 and BL2 – blood, SP1 and SP2 – sperm), NC sperm samples in teal, AMI sperm samples in orange and NMI sperm in purple. Box plots elements are defined as follows: center line: median; box limits: upper and lower quartiles; whiskers: 1.5× interquartile range; points: outliers. **Fig. S6:** Validation of *DDX4* and *XIST* methylation levels with deep bisulfite sequencing. A) Example of deep bisulfite sequencing results of *DDX4* and *XIST* in the three groups: NC, AMI and NMI. Each horizontal line of a plot represents a unique sequence read, while each vertical position represents a CpG site (methylated sites in red, unmethylated sites in blue). B) Mean methylation values for *DDX4* and *XIST* in the NC (teal, n=5), AMI (red, n=5) and NMI (purple, n=6) selected for the WGBS. **Fig. S7:** Deep bisulfite sequencing results of the AMI sample SOAT6 with atypical *H19* methylation pattern. **Fig. S8:** Deep bisulfite sequencing results for a representative NC sample**. Fig. S9:** Box plots showing for the 17 validated DMRs the distribution of the WGBS mean methylation values for the subset of CpGs covered by the targeted DBS approach (NC, teal, n = 5; NMI, purple, n = 6; Additional file 1: Table S9) and the targeted DBS methylation values for the validation NC samples (VNC, light teal, n = 20; Additional file 1: Table S11). Box plots elements are defined as follows: center line: median; box limits: upper and lower quartiles; whiskers: 1.5× interquartile range; points: outliers. **Fig. S10:** A) Principal component analysis (PCA) of the 14 CpG sites in the *H19* locus obtained by DBS for the 40 normal controls (NC, teal), 77 normally methylated infertile (NMI, purple) and one abnormally methylated infertile (AMI) (Additional file 1: Table S13). B) Contribution of the variables (14 CpG sites) to the principal components. (PDF 2,358 kb).

## References

1. Lazaraviciute G, Kauser M, Bhattacharya S, Haggarty P, Bhattacharya S: A systematic review and meta-analysis of DNA methylation levels and imprinting disorders in children conceived by IVF/ICSI compared with children conceived spontaneously. Hum Reprod Update 2014, 20(6):840–852.

2. Gosden R, Trasler J, Lucifero D, Faddy M: Rare congenital disorders, imprinted genes, and assisted reproductive technology. Lancet 2003, 361(9373):1975–1977.

3. Kläver R, Tuttelmann F, Bleiziffer A, Haaf T, Kliesch S, Gromoll J: DNA methylation in spermatozoa as a prospective marker in andrology. Andrology-Us 2013, 1(5):731–740.

4. Marques CJ, Carvalho F, Sousa M, Barros A: Genomic imprinting in disruptive spermatogenesis. Lancet 2004, 363(9422):1700–1702.

5. Poplinski A, Tuttelmann F, Kanber D, Horsthemke B, Gromoll J: Idiopathic male infertility is strongly associated with aberrant methylation of MEST and IGF2/H19 ICR1. Int J Androl 2010, 33(4):642–649.

6. Urdinguio RG, Bayon GF, Dmitrijeva M, Torano EG, Bravo C, Fraga MF, Bassas L, Larriba S, Fernandez AF: Aberrant DNA methylation patterns of spermatozoa in men with unexplained infertility. Hum Reprod 2015, 30(5):1014–1028.

7. Tang L, Liu Z, Zhang R, Su C, Yang W, Yao Y, Zhao S: Imprinting alterations in sperm may not significantly influence ART outcomes and imprinting patterns in the cord blood of offspring. PLoS One 2017, 12(11):e0187869.

8. Seisenberger S, Peat JR, Hore TA, Santos F, Dean W, Reik W: Reprogramming DNA methylation in the mammalian life cycle: building and breaking epigenetic barriers. Philos Trans R Soc Lond B Biol Sci 2013, 368(1609):20110330.

9. Seisenberger S, Andrews S, Krueger F, Arand J, Walter J, Santos F, Popp C, Thienpont B, Dean W, Reik W: The dynamics of genome-wide DNA methylation reprogramming in mouse primordial germ cells. Mol Cell 2012, 48(6):849–862.

10. Guo F, Yan L, Guo H, Li L, Hu B, Zhao Y, Yong J, Hu Y, Wang X, Wei Y et al: The Transcriptome and DNA Methylome Landscapes of Human Primordial Germ Cells. Cell 2015, 161(6):1437–1452.

11. Langenstroth-Röwer D, Gromoll J, Wistuba J, Trondle I, Laurentino S, Schlatt S, Neuhaus N: De novo methylation in male germ cells of the common marmoset monkey occurs during postnatal development and is maintained in vitro. Epigenetics 2017, 12(7):527–539.

12. Buiting K, Dittrich B, Gross S, Lich C, Farber C, Buchholz T, Smith E, Reis A, Burger J, Nothen MM et al: Sporadic imprinting defects in Prader-Willi syndrome and Angelman syndrome: implications for imprint-switch models, genetic counseling, and prenatal diagnosis. Am J Hum Genet 1998, 63(1):170–180.

13. Laurentino S, Beygo J, Nordhoff V, Kliesch S, Wistuba J, Borgmann J, Buiting K, Horsthemke B, Gromoll J: Epigenetic germline mosaicism in infertile men. Hum Mol Genet 2015, 24(5):1295–1304.

14. Ehmcke J, Schlatt S: A revised model for spermatogonial expansion in man: lessons from non-human primates. Reproduction 2006, 132(5):673–680.

15. Ehmcke J, Wistuba J, Schlatt S: Spermatogonial stem cells: questions, models and perspectives. Hum Reprod Update 2006, 12(3):275–282.

16. Santi D, De Vincentis S, Magnani E, Spaggiari G: Impairment of sperm DNA methylation in male infertility: a meta-analytic study. Andrology-Us 2017, 5(4):695–703.

17. Marques CJ, Costa P, Vaz B, Carvalho F, Fernandes S, Barros A, Sousa M: Abnormal methylation of imprinted genes in human sperm is associated with oligozoospermia. Molecular Human Reproduction 2008, 14(2):67–73.

18. Laqqan M, Solomayer EF, Hammadeh M: Aberrations in sperm DNA methylation patterns are associated with abnormalities in semen parameters of subfertile males. Reprod Biol 2017, 17(3):246–251.

19. Ziller MJ, Hansen KD, Meissner A, Aryee MJ: Coverage recommendations for methylation analysis by whole-genome bisulfite sequencing. Nat Methods 2015, 12(3):230–232, 231 p following 232.

20. Laurentino S, Cremers J-F, Horsthemke B, Tuettelmann F, Czeloth K, Zitzmann M, Pohl E, Rahmann S, Schroeder C, Berres S et al: Healthy ageing men have normal reproductive function but display germline-specific molecular changes. 2019:19006221.

21. Monk D, Morales J, den Dunnen JT, Russo S, Court F, Prawitt D, Eggermann T, Beygo J, Buiting K, Tumer Z et al: Recommendations for a nomenclature system for reporting methylation aberrations in imprinted domains. Epigenetics 2018, 13(2):117–121.

22. Hermann BP, Cheng K, Singh A, Roa-De La Cruz L, Mutoji KN, Chen IC, Gildersleeve H, Lehle JD, Mayo M, Westernstroer B et al: The Mammalian Spermatogenesis Single-Cell Transcriptome, from Spermatogonial Stem Cells to Spermatids. Cell Rep 2018, 25(6):1650–1667 e1658.

23. Luján S, Caroppo E, Niederberger C, Arce JC, Sadler-Riggleman I, Beck D, Nilsson E, Skinner MK: Sperm DNA Methylation Epimutation Biomarkers for Male Infertility and FSH Therapeutic Responsiveness. Sci Rep 2019, 9(1):16786.

24. Laurentino S, Heckmann L, Di Persio S, Li X, Meyer Zu Horste G, Wistuba J, Cremers JF, Gromoll J, Kliesch S, Schlatt S et al: High-resolution analysis of germ cells from men with sex chromosomal aneuploidies reveals normal transcriptome but impaired imprinting. Clin Epigenetics 2019, 11(1):127.

25. Do C, Shearer A, Suzuki M, Terry MB, Gelernter J, Greally JM, Tycko B: Genetic-epigenetic interactions in cis: a major focus in the post-GWAS era. Genome Biol 2017, 18(1):120.

26. Boissonnas CC, El Abdalaoui H, Haelewyn V, Fauque P, Dupont JM, Gut I, Vaiman D, Jouannet P, Tost J, Jammes H: Specific epigenetic alterations of IGF2-H19 locus in spermatozoa from infertile men. Eur J Hum Genet 2010, 18(1):73–80.

27. Camprubi C, Pladevall M, Grossmann M, Garrido N, Pons MC, Blanco J: Semen samples showing an increased rate of spermatozoa with imprinting errors have a negligible effect in the outcome of assisted reproduction techniques. Epigenetics 2012, 7(10):1115–1124.

28. Dong H, Wang YX, Zou ZK, Chen LM, Shen CY, Xu SQ, Zhang J, Zhao FF, Ge SQ, Gao Q et al: Abnormal Methylation of Imprinted Genes and Cigarette Smoking: Assessment of Their Association With the Risk of Male Infertility. Reprod Sci 2017, 24(1):114–123.

29. Li B, Li JB, Xiao XF, Ma YF, Wang J, Liang XX, Zhao HX, Jiang F, Yao YQ, Wang XH: Altered DNA methylation patterns of the H19 differentially methylated region and the DAZL gene promoter are associated with defective human sperm. PLoS One 2013, 8(8):e71215.

30. Li XP, Hao CL, Wang Q, Yi XM, Jiang ZS: H19 gene methylation status is associated with male infertility. Experimental and Therapeutic Medicine 2016, 12(1):451–456.

31. Marques PI, Fernandes S, Carvalho F, Barros A, Sousa M, Marques CJ: DNA methylation imprinting errors in spermatogenic cells from maturation arrest azoospermic patients. Andrology-Us 2017, 5(3):451–459.

32. Minor A, Chow V, Ma S: Aberrant DNA methylation at imprinted genes in testicular sperm retrieved from men with obstructive azoospermia and undergoing vasectomy reversal. Reproduction 2011, 141(6):749–757.

33. Ward LD, Kellis M: HaploReg v4: systematic mining of putative causal variants, cell types, regulators and target genes for human complex traits and disease. Nucleic Acids Res 2016, 44(D1):D877–881.

34. Coolen MW, Statham AL, Qu WJ, Campbell MJ, Henders AK, Montgomery GW, Martin NG, Clark SJ: Impact of the Genome on the Epigenome Is Manifested in DNA Methylation Patterns of Imprinted Regions in Monozygotic and Dizygotic Twins. Plos One 2011, 6(10).

35. Marjonen H, Kahila H, Kaminen-Ahola N: rs1073251 IGF2/H19 locus associates with a genotype-specific trend in placental DNA methylation and head circumference of prenatally alcohol-exposed newborns. Hum Reprod Open 2017, 2017(3):hox014.

36. Kläver R, Gromoll J: Bringing epigenetics into the diagnostics of the andrology laboratory: challenges and perspectives. Asian J Androl 2014, 16(5):669–674.

37. Gambera L, Serafini F, Morgante G, Focarelli R, De Leo V, Piomboni P: Sperm quality and pregnancy rate after COX-2 inhibitor therapy of infertile males with abacterial leukocytospermia. Hum Reprod 2007, 22(4):1047–1051.

38. Jenkins TG, Liu L, Aston KI, Carrell DT: Pre-screening method for somatic cell contamination in human sperm epigenetic studies. Syst Biol Reprod Med 2018, 64(2):146–155.

39. Horsthemke B: In brief: genomic imprinting and imprinting diseases. J Pathol 2014, 232(5):485–487.

40. Mackay DJG, Temple IK: Human imprinting disorders: Principles, practice, problems and progress. Eur J Med Genet 2017, 60(11):618–626.

41. World Health Organization.: WHO laboratory manual for the examination and processing of human semen, 5th edn. Geneva: World Health Organization; 2010.

42. Souren NY, Gerdes LA, Lutsik P, Gasparoni G, Beltrán E, Salhab A, Kümpfel T, Weichenhan D, Plass C, Hohlfeld R et al: DNA methylation signatures of monozygotic twins clinically discordant for multiple sclerosis. Nature Communications 2019, 10(1):2094.

43. Pedersen BS, Eyring K, De S, Yang IV, Schwartz DA: Fast and accurate alignment of long bisulfite-seq reads. arXiv 2014, 1401.1129 [q-bio.GN].

44. Picard toolkit. Broad Institute [https://github.com/broadinstitute/picard]

45. MethylDackel [https://github.com/dpryan79/MethylDackel]

46. Ewels P, Magnusson M, Lundin S, Kaller M: MultiQC: summarize analysis results for multiple tools and samples in a single report. Bioinformatics 2016, 32(19):3047–3048.

47. FastQC [http://www.bioinformatics.babraham.ac.uk/projects/fastqc]

48. Okonechnikov K, Conesa A, Garcia-Alcalde F: Qualimap 2: advanced multi-sample quality control for high-throughput sequencing data. Bioinformatics 2016, 32(2):292–294.

49. Schröder C: Bioinformatics from genetic variants to methylation. 2018.

50. Juhling F, Kretzmer H, Bernhart SH, Otto C, Stadler PF, Hoffmann S: metilene: fast and sensitive calling of differentially methylated regions from bisulfite sequencing data. Genome Res 2016, 26(2):256–262.

51. Pedersen BS, Quinlan AR: Mosdepth: quick coverage calculation for genomes and exomes. Bioinformatics 2018, 34(5):867–868.

52. Lawrence M, Huber W, Pages H, Aboyoun P, Carlson M, Gentleman R, Morgan MT, Carey VJ: Software for computing and annotating genomic ranges. PLoS Comput Biol 2013, 9(8):e1003118.

53. Haeussler M, Zweig AS, Tyner C, Speir ML, Rosenbloom KR, Raney BJ, Lee CM, Lee BT, Hinrichs AS, Gonzalez JN et al: The UCSC Genome Browser database: 2019 update. Nucleic Acids Res 2019, 47(D1):D853–D858.

54. Cunningham F, Achuthan P, Akanni W, Allen J, Amode MR, Armean IM, Bennett R, Bhai J, Billis K, Boddu S et al: Ensembl 2019. Nucleic Acids Res 2019, 47(D1):D745–D751.

55. Leitão E, Beygo J, Zeschnigk M, Klein-Hitpass L, Bargull M, Rahmann S, Horsthemke B: Locus-Specific DNA Methylation Analysis by Targeted Deep Bisulfite Sequencing. Methods Mol Biol 2018, 1767:351–366.

56. R: A language and environment for statistical computing [https://www.R-project.org/]

