## Additional file 2 for "The sperm epigenome does not display recurrent epimutations in patients with severely impaired spermatogenesis"

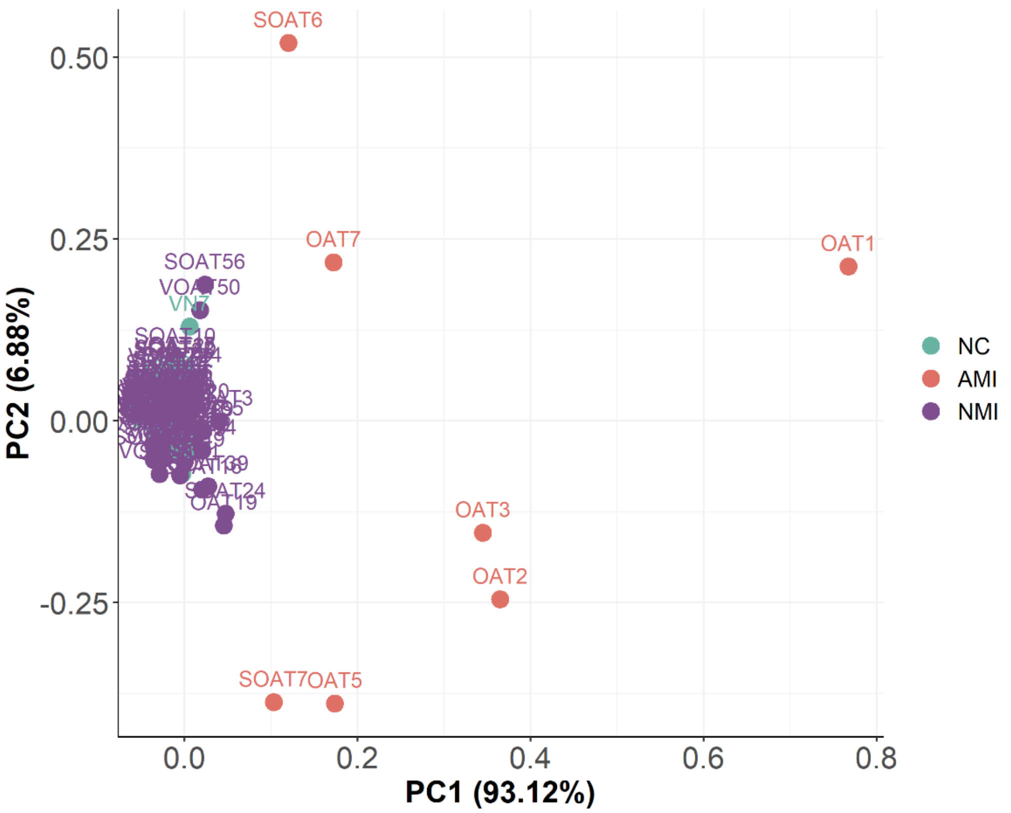


Fig. S1: Principle component analysis (PCA) of *MEST* and *H19* methylation values obtained by DBS for the 133 sperm samples (Additional file 1: Table S2). Samples with PC1 score < 0.1 were considered normally methylated and with PC1 score ≥ 0.1, abnormally methylated. While normal controls (NC) are a homogeneous group of normally methylated samples, infertile sperm samples were subdivided in two groups according to this PC1 threshold (AMI, abnormally methylated infertile; NMI, normally methylated infertile).

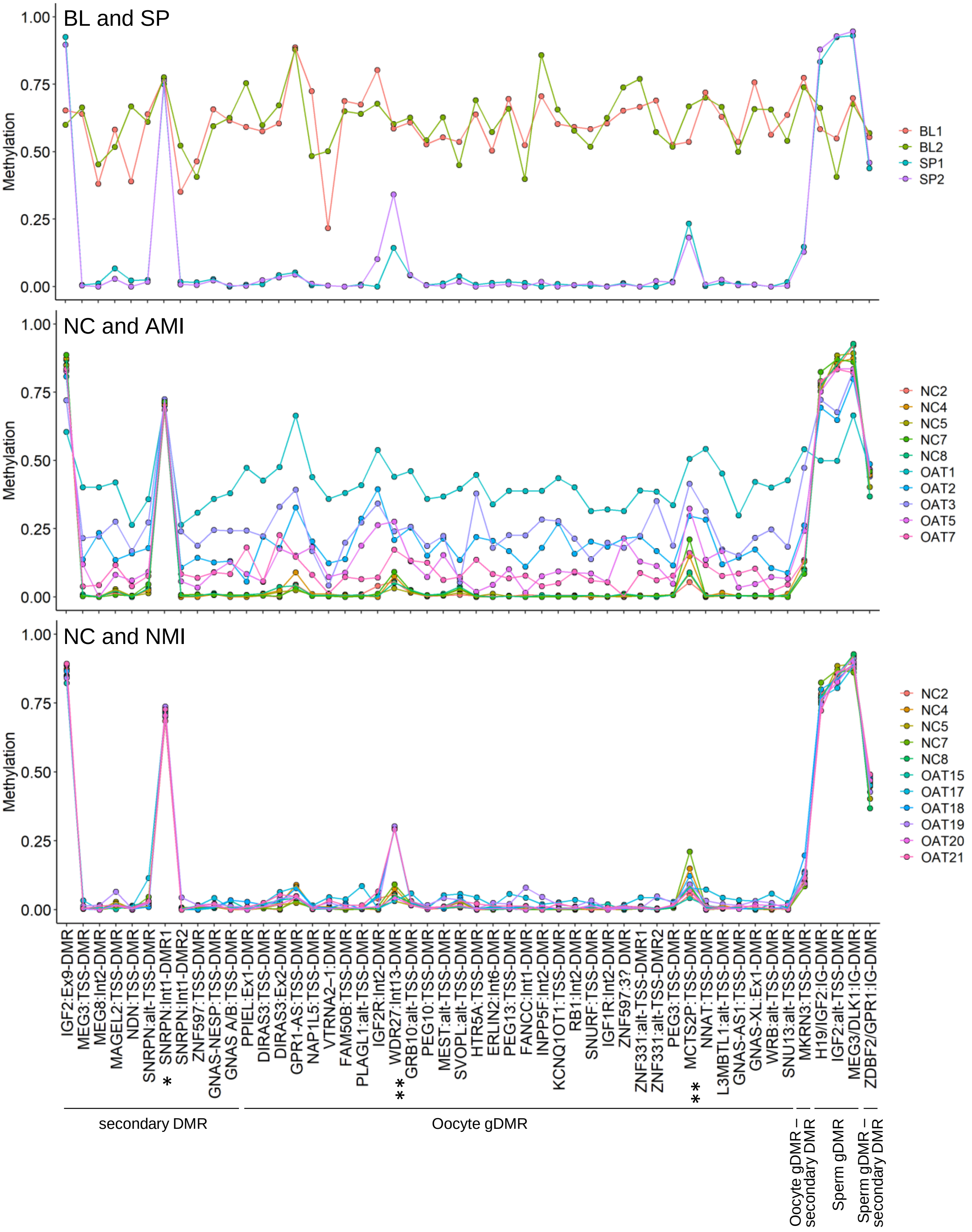


Fig. S2: Methylation levels of the 50 imprinting control regions. Line diagrams showing comparisons between blood (BL1, BL2) and sperm (SP1, SP2) datasets from Laurentino et al. [20] (upper panel), between NC and AMI sperm (middle) and NC and AMI sperm (lower panel) (Additional file 1: Table S4). * Not imprinted according to this data, ** Possible polymorphism.

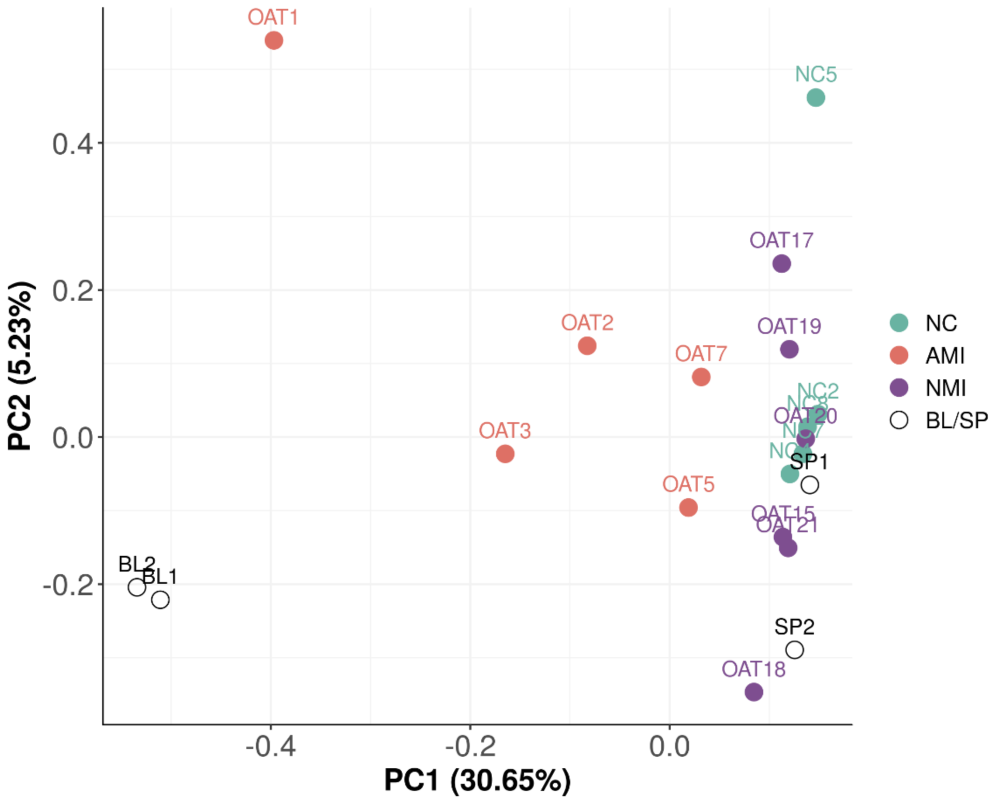


Fig. S3: PCA generated for ~8.7 million CpG loci where all samples show methylation values. Only loci with minimum coverage of five in all samples and minimum mapping quality of 10 are considered. Datasets from Laurentino et al. [20] are shown in white (BL1 and BL2 – blood, SP1 and SP2 – sperm), NC sperm samples in teal, AMI sperm samples in orange and NMI sperm in purple.

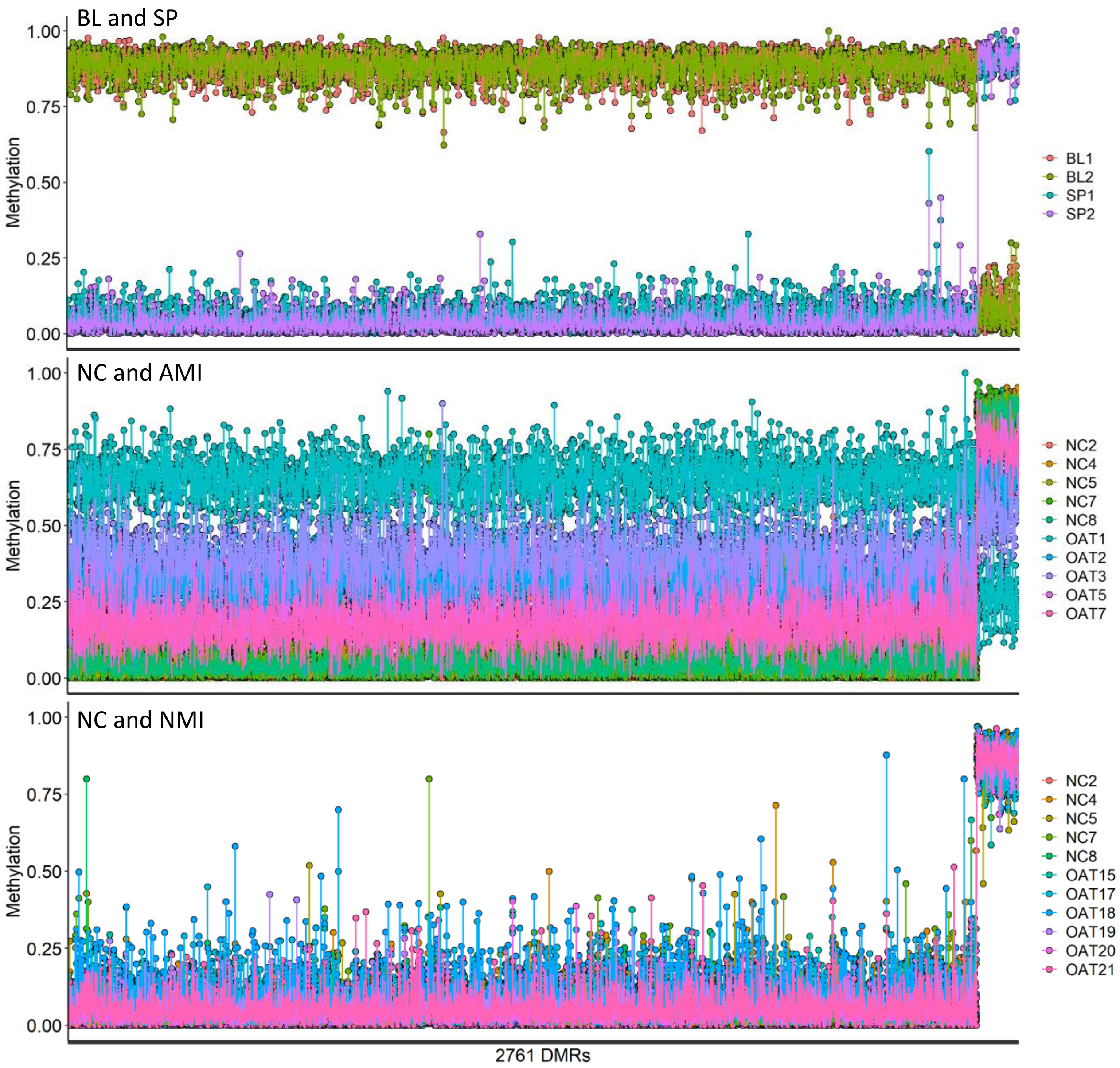


Fig. S4: Methylation levels of 2,761 sperm-soma DMRs. Line diagrams showing comparisons between blood (BL1, BL2) and sperm (SP1, SP2) datasets from Laurentino et al. [20] (upper panel), between NC and AMI sperm (middle) and NC and NMI sperm (lower panel). The 2,640 DMRs less methylated in sperm than in blood are towards the left and the 121 DMRs more methylated in sperm than in blood are on the right (Additional file 1: Table S5).

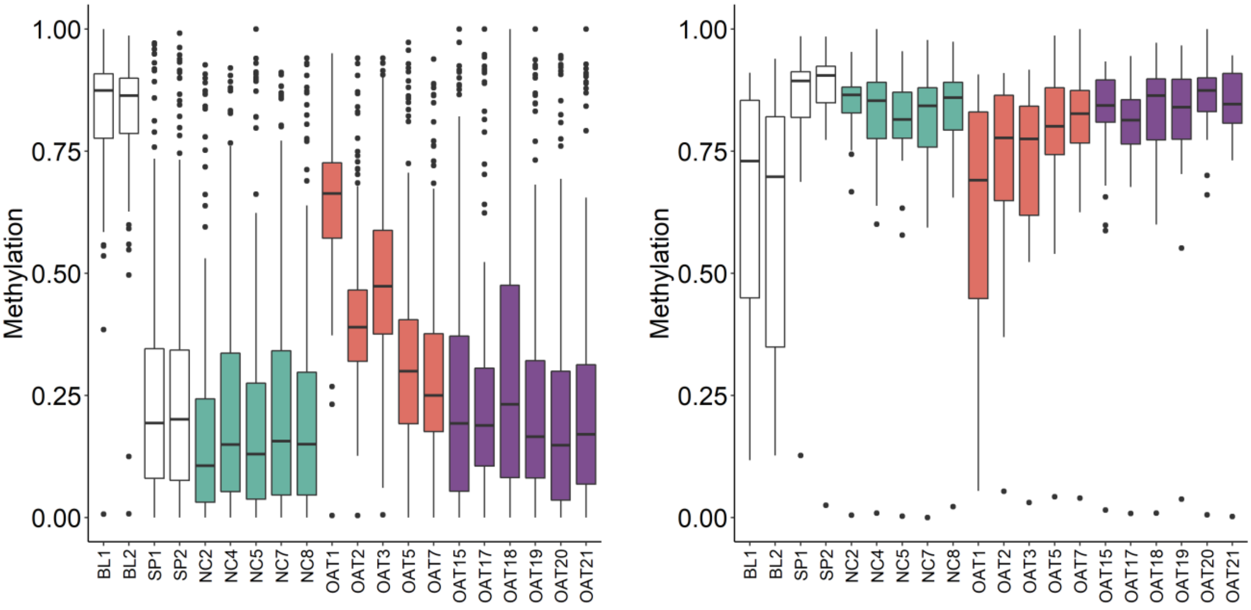


Fig. S5: Methylation levels of the 217 DMRs claimed by Luján et al. [23] to be useful for infertility assessment. Box plots showing the distribution of methylation values for the DMRs stated to be hyper- (190 DMRs, left) or hypomethylated in sperm from infertile vs. fertile men (20 DMRs, right) (Additional file 1: Table S6). Datasets from Laurentino et al. [20] are shown in white (BL1 and BL2 – blood, SP1 and SP2 – sperm), NC sperm samples in teal, AMI sperm samples in orange and NMI sperm in purple. Box plots elements are defined as follows: center line: median; box limits: upper and lower quartiles; whiskers: 1.5× interquartile range; points: outliers.

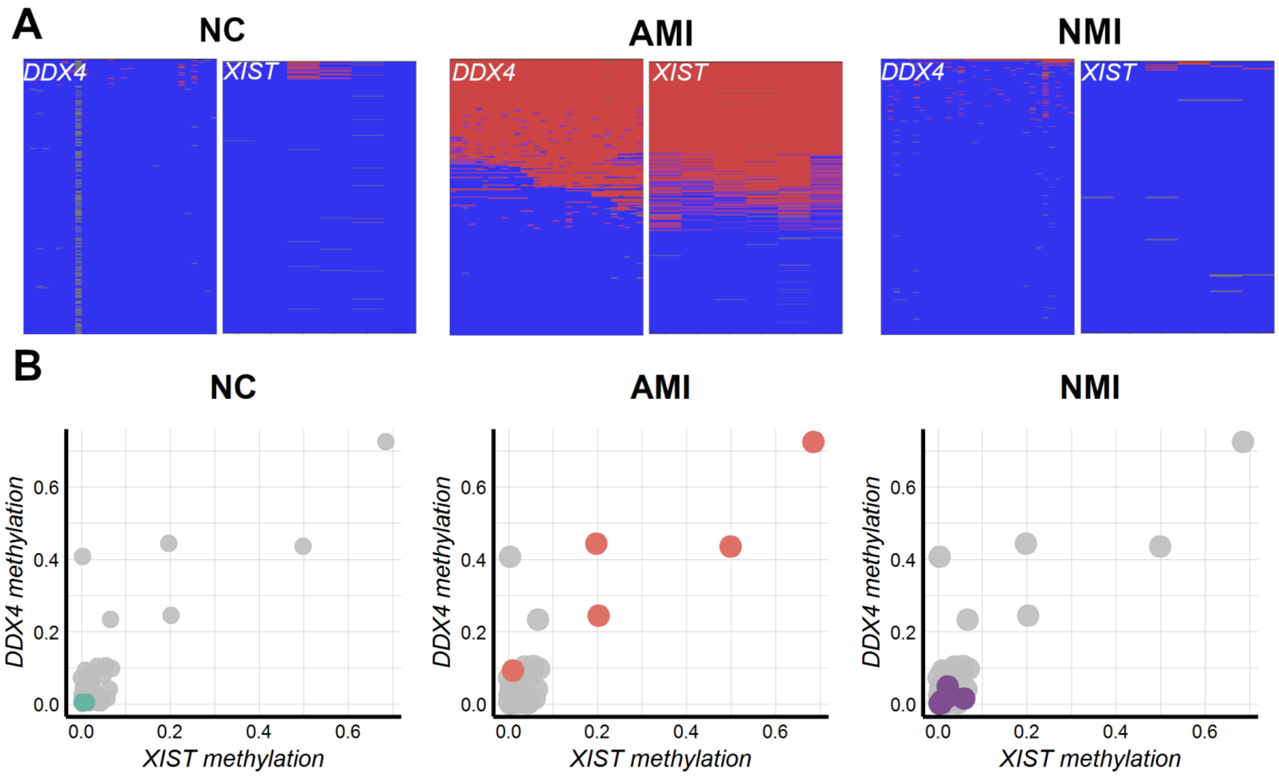


Fig. S6: Validation of *DDX4* and *XIST* methylation levels with deep bisulfite sequencing. A) Example of deep bisulfite sequencing results of *DDX4* and *XIST* in the three groups: NC, AMI and NMI. Each horizontal line of a plot represents a unique sequence read, while each vertical position represents a CpG site (methylated sites in red, unmethylated sites in blue). B) Mean methylation values for *DDX4* and *XIST* in the NC (teal, n=5), AMI (red, n=5) and NMI (purple, n=6) selected for the WGBS.

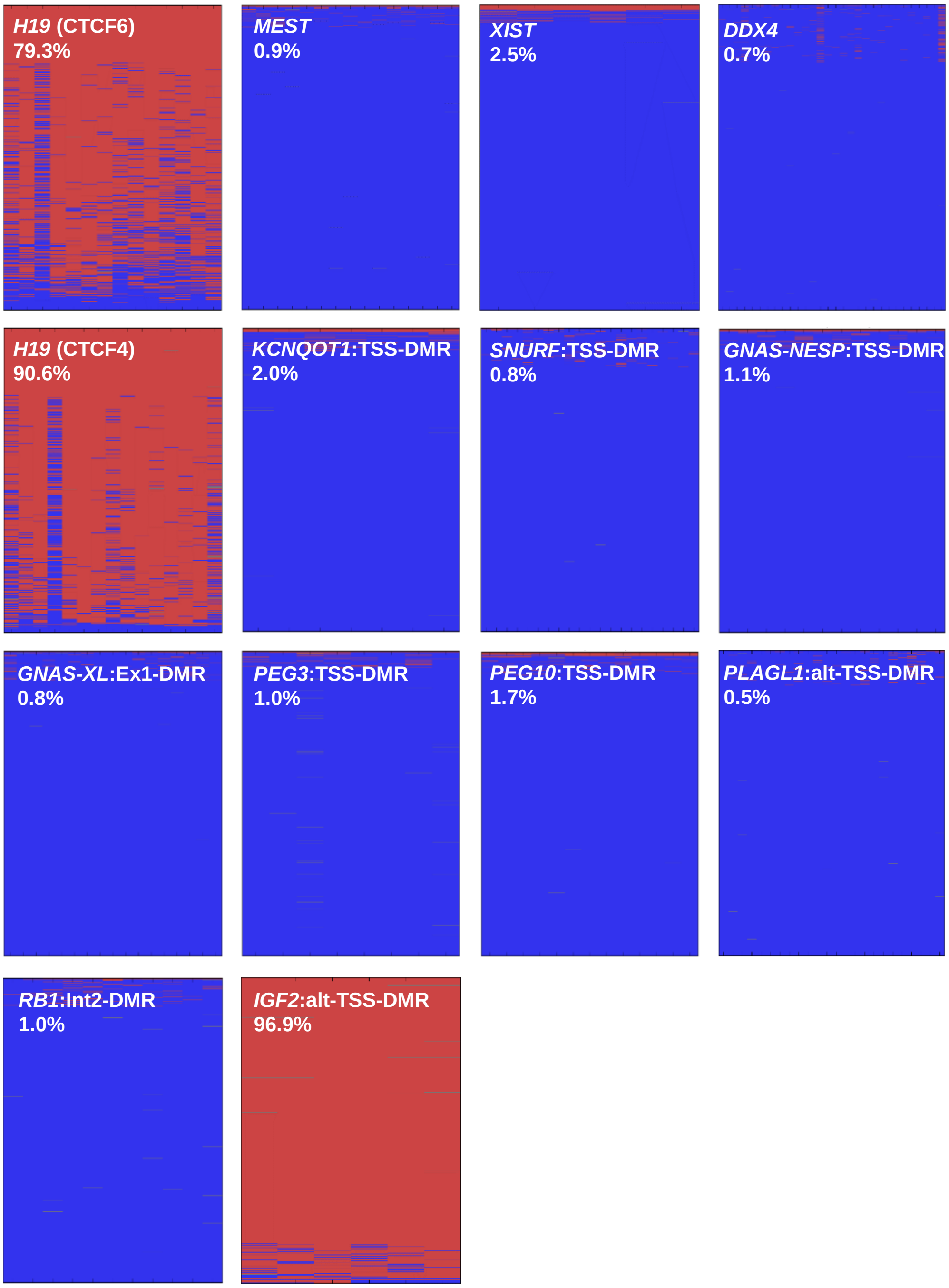


Fig. S7: Deep bisulfite sequencing results of the AMI sample SOAT6 with atypical *H19* methylation pattern.

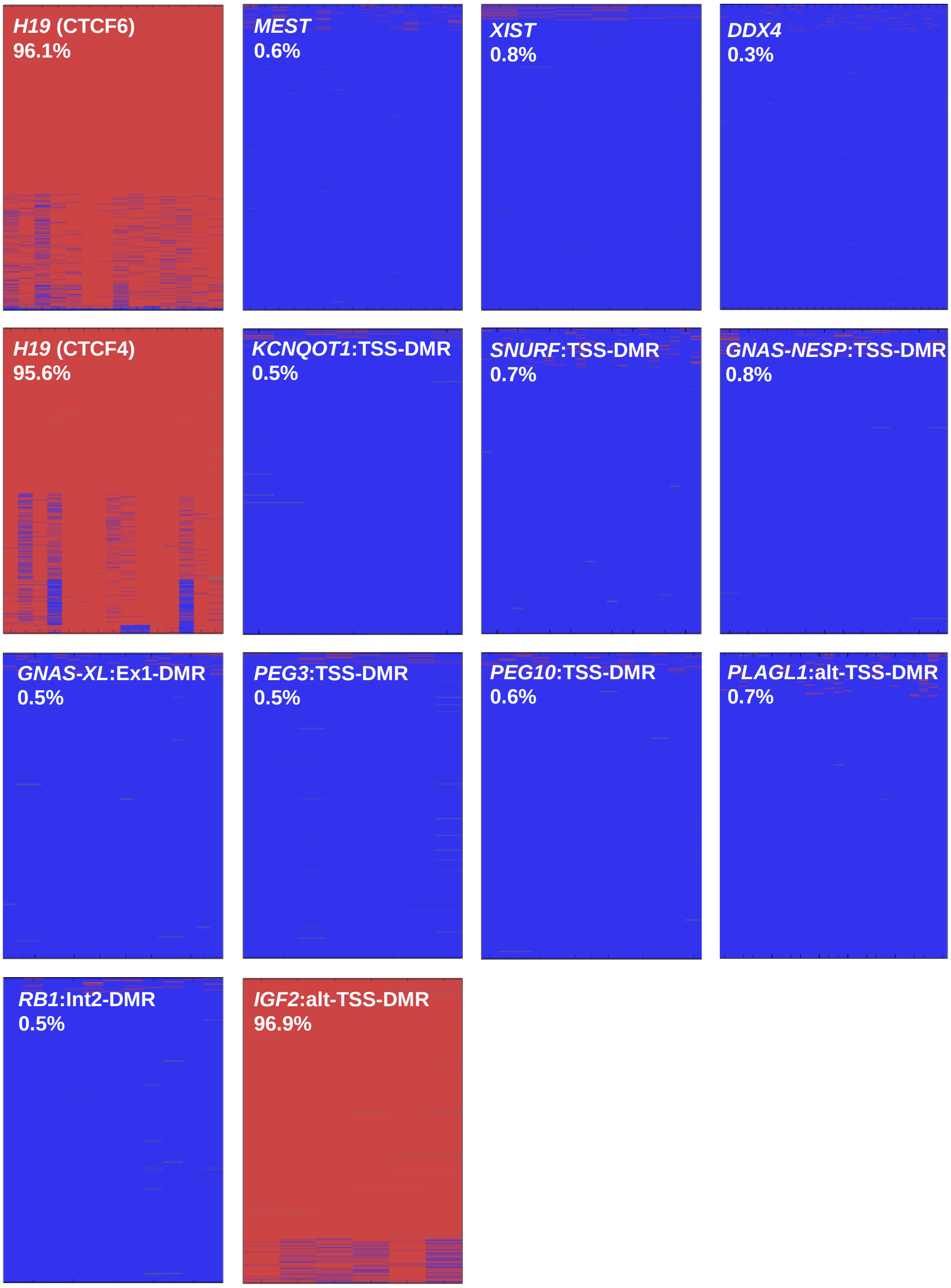


Fig. S8: Deep bisulfite sequencing results for a representative NC sample.

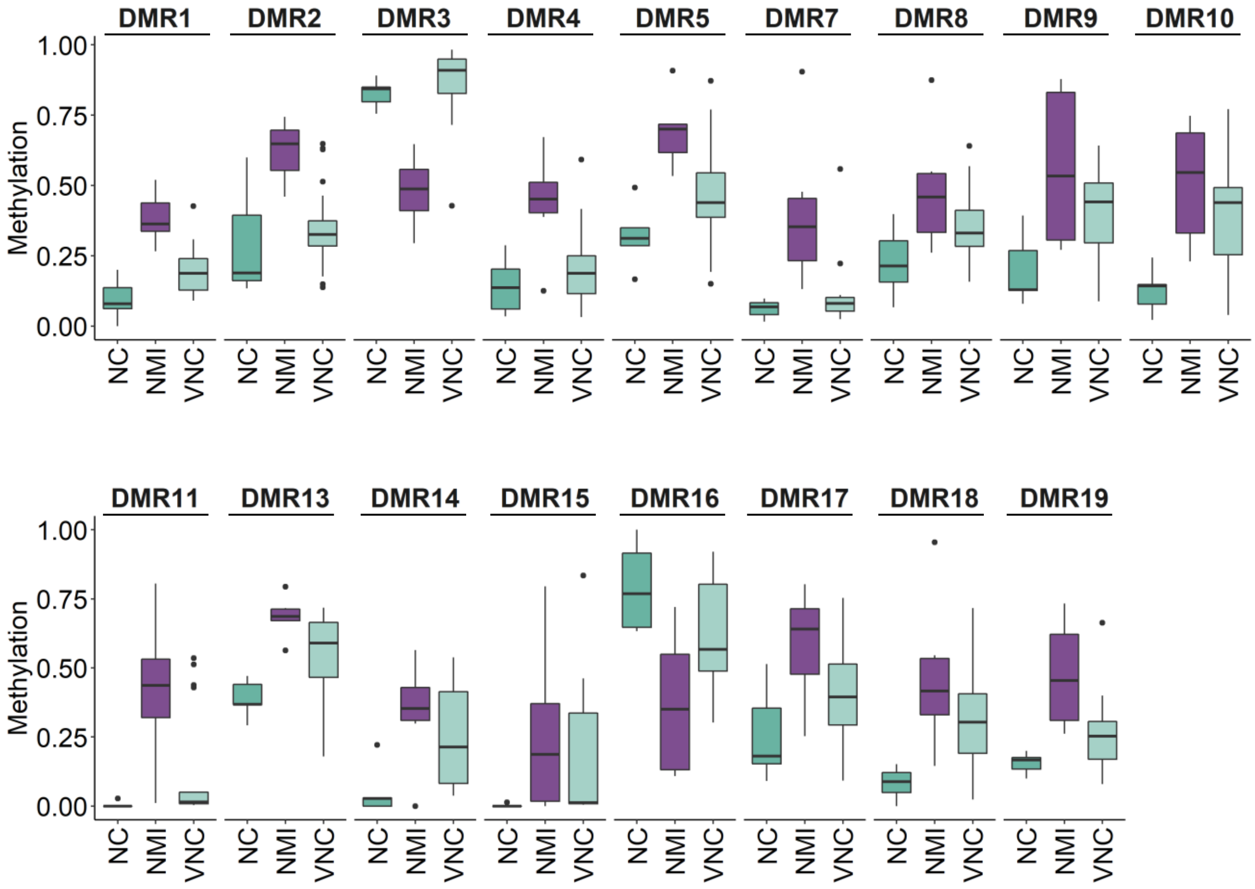


Fig. S9: Box plots showing for the 17 validated DMRs the distribution of the WGBS mean methylation values for the subset of CpGs covered by the targeted DBS approach (NC, teal, n = 5; NMI, purple, n = 6; Additional file 1: Table S9) and the targeted DBS methylation values for the validation NC samples (VNC, light teal, n = 20; Additional file 1: Table S11). Box plots elements are defined as follows: center line: median; box limits: upper and lower quartiles; whiskers: 1.5× interquartile range; points: outliers.

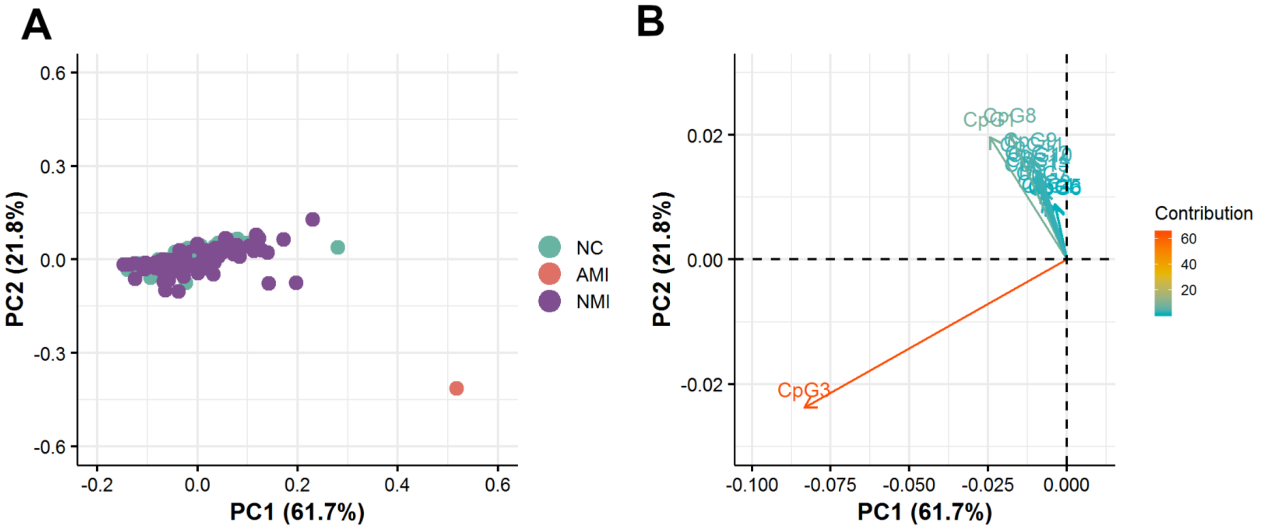


Fig. S10: A) Principal component analysis (PCA) of the 14 CpG sites in the *H19* locus obtained by DBS for the 40 normal controls (NC, teal), 77 normally methylated infertile (NMI, purple) and one abnormally methylated infertile (AMI) (Additional file 1: Table S13). B) Contribution of the variables (14 CpG sites) to the principal components.
